# Correlation structure of grid cells is preserved during sleep

**DOI:** 10.1101/198499

**Authors:** Richard J. Gardner, Li Lu, Tanja Wernle, May-Britt Moser, Edvard I. Moser

**Affiliations:** Kavli Institute for Systems Neuroscience and Centre for Neural Computation, Norwegian University of Science and Technology, Trondheim, Norway

**Author notes:** Corresponding author: Richard J. Gardner; Edvard I. Moser. Present address: Li Lu: Department of Neuroscience, Baylor College of Medicine, Houston, USA. Tanja Wernle: Friedrich Miescher Institute for Biomedical Research, Maulbeerstrasse 66, CH-4058 Basel, Switzerland. **Author contributions**: R.J.G., M.-B.M. and E.I.M. designed the experiment, R.J.G. collected most data but used additional recordings from L.L. and T.W., R.J.G. analyzed the data, and R.J.G. wrote the paper, assisted by M.-B.M. and E.I.M. All authors read and commented on the paper. M.-B.M. and E.I.M. supervised the project.

## Abstract

The network of grid cells in the medial entorhinal cortex forms a fixed reference frame for mapping physical space. The mechanistic origin of the grid representation is unknown, but continuous attractor network (CAN) models explain multiple fundamental features of grid-cell activity. An untested prediction of CAN grid models is that the grid-cell network should exhibit an activity correlation structure that transcends behavioural or brain states. By recording from MEC cell ensembles during navigation and sleep, we found that spatial phase offsets of grid cells predict arousal-state-independent spike rate correlations. Similarly, state-invariant correlations between conjunctive grid-head-direction and pure head-direction cells were predicted by their head-direction tuning offsets. Spike rates of grid cells were only weakly correlated across modules, and module scale relationships disintegrated during slow-save sleep, suggesting that modules function as independent attractor networks. Collectively, our observations suggest that network states in MEC are expressed universally across brain and behaviour states.

---

A fundamental goal in systems neuroscience is to uncover the mechanistic and structural bases for information processing in the brain *in vivo*. The sensory pathways of the mammalian brain have over decades become established as model systems for studying this matter because they endow experimenters with the power to precisely and systematically manipulate stimuli while observing neuronal responses^1,2^. Additionally, neurons in primary sensory cortex typically encode low-dimensional stimulus features^3,4^, making it possible to draw clear relationships between input signals and neuronal responses and thus deduce the neural code.

In higher-order sensory and association cortex, our understanding of coding and computation is comparatively poor. Activity within these areas comprises a complex integration of many signals, making interpretation and experimental dissection of the encoded information highly challenging. A notable exception to this rule is the mammalian brain’s “cognitive map”, which employs a system of neurons that represents an animal’s location allocentrically within its physical environment^5,6^. Crucially, this map is not dependent on or responsive to any particular sensory modality, but instead appears to be a high-order cognitive abstraction based on multimodal sensory cues and self-motion information^7,8^. Several functionally and anatomically distinct classes of neurons in the hippocampal region are part of the cognitive map – notably place cells, head-direction (HD) cells, border cells, and grid cells, each of which signals a different aspect of the animal’s spatial location in its environment^9–13^.

The overarching question of how such robust low-dimensional neuronal representations are maintained in a highly dynamic sensory and behavioural context has attracted widespread attention from both experimentalists and theoreticians. In relation to grid cells, a variety of computational network models have demonstrated that in principle, there are multiple distinct mechanisms by which periodic grid patterns might be generated^6,14^. One group of models is based on continuous attractor network (CAN) dynamics: a phenomenon in which network activity reliably converges on an intrinsic manifold consisting of a continuum of equally favourable states, allowing a network activity pattern to be smoothly translated across the manifold^15^. CAN models have gained credence in the field because they predict several essential properties of place cells and grid cells^6–8,16–19^. Specifically, the models predict that grid cells within a functional network should share a common spacing and orientation, and that grid cells should have fixed grid phase (translational offset) relationships across different environments^6,8^. A corollary of these predictions is that independent grid spacing, orientation, or spatial phase offsets could only exist through a modular organization of functionally discrete grid networks^8,20^. Indeed, all of these predictions have been experimentally confirmed^21–24^. However, it remains uninvestigated whether these signatures of CAN dynamics are expressed outside of the active explorative state.

An important feature of all attractor network models is that the grid phase offset of any two grid cells in the network is a function of the net synaptic connectivity between them (the cells with the highest net connectivity have the smallest spatial phase offsets). Thus, the network’s design prescribes an intrinsic correlation structure which should manifest as “noise” correlation, irrespective of the network’s mode of activity. Models without continuous attractor characteristics do not, in general, make this prediction – therefore, probing for this specific correlation structure is an assay for the mechanisms described in grid CAN models.

An attractive experimental approach for measuring the intrinsic correlation structure of the grid cell network would be to record grid cell activity when the spatial signal is absent and the network is instead predominated by self-organized dynamics. During such a state, correlations observed in the grid cell network should be a close reflection of the circuit’s synaptic organization. An ideal candidate for such a state is slow-wave sleep (SWS), during which afferent transmission of sensory information is suppressed, top-down cognitive influences are absent, and functional connectivity is more localized^25^, allowing internally generated brain dynamics to predominate. Indeed, in the rodent head-direction circuit, which has also been postulated to depend on continuous attractor mechanisms^26–28^, population activity remains constrained within the same low-dimensional state space across arousal states^29^. We therefore find it pertinent to ask whether the same could be true for grid cells.

We thus recorded from grid, HD and conjunctive grid-HD cell ensembles in the medial entorhinal cortex (MEC) and parasubiculum (PaS), while rats ran in an open field environment and also while they slept. We show that the grid phase relationships between intramodular grid-cell pairs during wakeful navigation are rigidly preserved during SWS as well as REM, in the form of temporal spiking correlations, consistent with the predictions of CAN models.

## Results

We obtained *in vivo* extracellular electrophysiological recordings from MEC and PaS in 7 rats, during foraging behaviour in a 1.5-metre square open field arena (RUN), as well as during natural sleep in a rest chamber, from which we extracted periods of slow-wave sleep (SWS) and rapid-eye-movement sleep (REM). Periods of REM and SWS were identified from sustained immobility, accompanied by signature patterns of activity in the MEC local field potential (LFP): SWS was signified by high-amplitude 1–4 Hz waves, while 5–10 Hz theta waves predominated during REM (**Supplementary Fig. 1a,b**). MEC or PaS single units were classified with established methods according to their spatial tuning properties during the RUN session (see methods). Units were identified as pure grid cells (*n* = 138), pure head-direction cells (HD, *n* = 95), or conjunctive grid × head-direction cells (grid-HD, *n* = 39).

Since the basic spiking properties of spatially tuned cells in MEC and PaS during sleep have not previously been described in detail, we first investigated whether these cells remained active during the states of SWS and REM. We fitted a repeated-measures ANOVA to the mean spike rates of the units in each state, with neuronal class as the between-subjects factor and arousal state as the within-subject factor. During each state, the mean spike rates of all three neuronal classes were similar (between-subjects effect of cell class *F*(2, 269) = 2.89, *P =* 0.0572; all post-hoc comparisons *P* > 0.05; Tukey-Kramer**; Supplementary Fig. 1c**). Individual cells of each class exhibited significant changes in mean rate between the three states, with rates lowest in SWS and highest in REM (within-subject effect of state *F*(2, 538) = 44.3, *P =* 1.57 × 10^−18^; all post-hoc state comparisons *P* < 0.01; **Supplementary Fig. 1c**). RUN, SWS and REM were accompanied by markedly different patterns of synchronous population activity (**Supplementary Fig. 1a,b**). During RUN and REM, prominent theta oscillations were present in the LFP, and individual grid cells showed periods of elevated spiking in the order of seconds. During SWS, the LFP became irregular and was dominated by <5 Hz waves accompanied by large fluctuations in the population spike rate^30^.

We then set out to test the prediction of CAN models that the same low-dimensional network states are expressed across all brain behavioural states. This should manifest as a fixed structure of spiking correlations between pairs of grid cells tightly linked to the degree of correspondence in their spatial activation patterns. Specifically, grid cells with overlapping or nonoverlapping spatial fields should respectively exhibit positive and negative spiking correlations.

Spiking correlations represent the sum of all influences on a cell pair’s spiking. One notably strong influence on correlations can be the global population rate^31^. To allow us to isolate coupled spiking of units irrespective of their modulation by the population rate, we calculated pairwise couplings by fitting a simple Poisson generalized linear model (GLM) to the spike rate of one neuron (see methods). The model had two regressors: first the spike rate of the paired neuron, and second the population spike rate. We demonstrated with simulated spike trains that the GLM method was able to faithfully extract coupling between units where the Pearson correlation became distorted by population rate effects (**Supplementary Fig. 2a**).

We hence used the model’s coefficient for the coupling between the two cells’ rates, hereafter referred to as “*β*”, as our principal correlation metric. We calculated cross-correlations by iteratively applying the GLM analysis over a range of time lags between the two neurons’ spike trains. When applying the GLM to our single-unit recordings, we observed that units in each neuronal class were overwhelmingly positively coupled to the population rate (*P* < 10^−10^, *z* > 6 in all states, binomial sign test, *n* = 138 grid cells, *n* = 95 HD cells, *n =* 39 grid-HD cells; **Supplementary Fig. 2c**). Accordingly, we found that the GLM cell-pair couplings were negatively compensated with respect to the corresponding Pearson correlation *r*-values (**Supplementary Fig. 2b**).

We then identified pairs of co-recorded grid cells and quantified their phase offsets. We only considered pairs of grid cells that were recorded on different tetrodes, to avoid the potential confounds of spike sorting errors or missed detection of temporally overlapping spikes. Using a previously described method^32^ (**Fig. 2a–c**; see methods), we classified co-recorded grid-cell pairs as being from the same module (‘intramodular’, *n* = 135) or separate modules (‘transmodular’, *n* = 63). For each intramodular grid-cell pair, we calculated *ϕ*_*G*_, the magnitude of the grid phase offset vector during RUN. We also calculated spike rate cross-correlograms (SRCs) separately for RUN, SWS and REM epochs. Both the raw SRC histograms and the GLM SRCs of individual cell pairs were typically characterized by a single positive or negative peak close to zero-lag (**Fig. 1a, Supplementary Fig. 3a,b**). The polarity and size of the zero-lag peak was generally conserved between states, although the SWS peaks were markedly compressed in time relative to RUN and REM. Cell pairs with the largest phase offsets showed near mutual exclusivity of spiking at zero-lag (**Fig. 1a2**).

**Figure 1:**
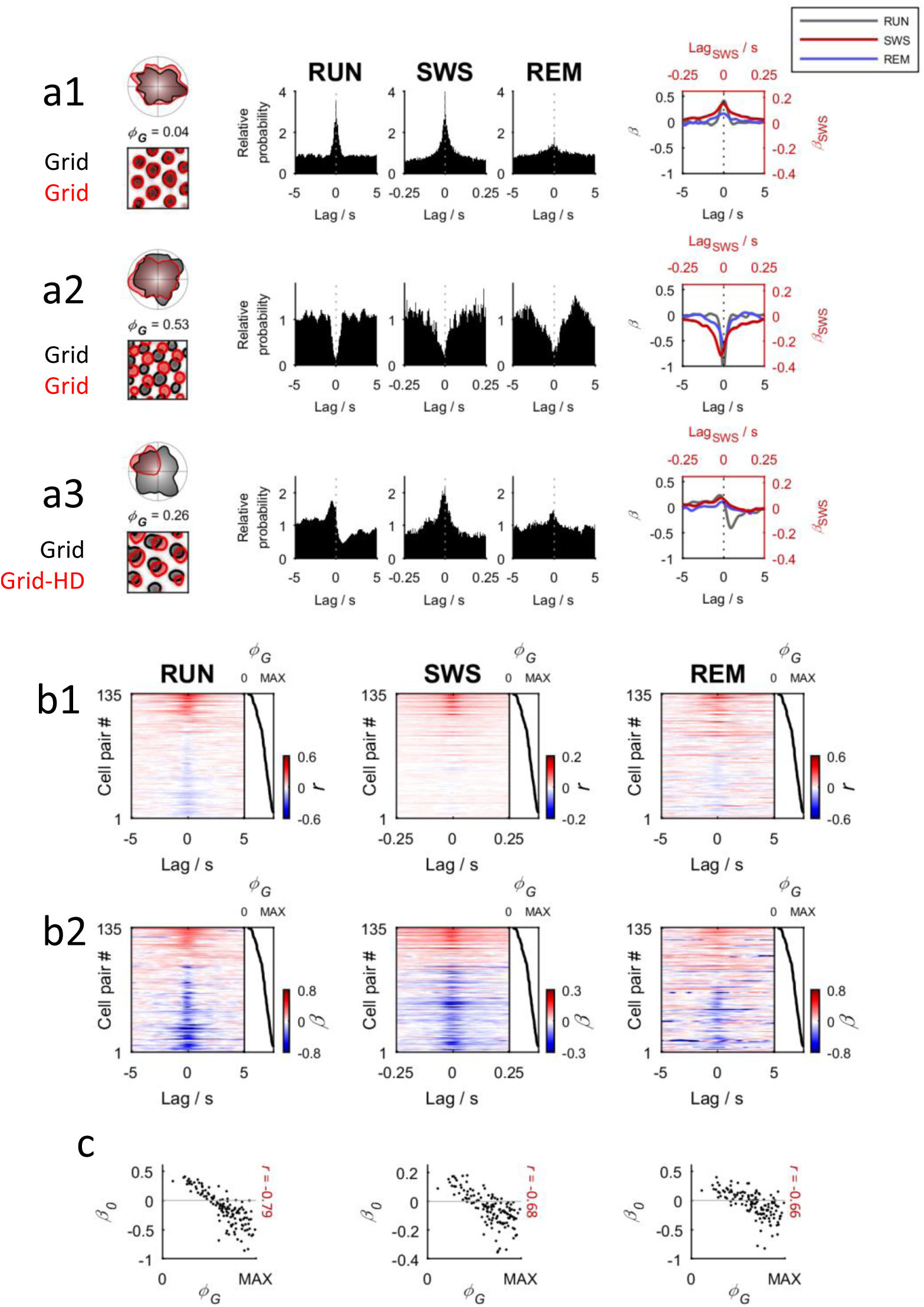
Grid spatial phase and temporal spiking correlations. **a:** Grid phase-offset and spike-timing relationships of two representative grid/grid cell pairs (rows **a1** and **a2**) and a grid/grid-HD cell pair (row **a3**). Top left of each row: head-direction tuning curve of both units of the cell pair during RUN, one in red shade and one in grey. Bottom left of each row: superimposed 2D firing rate maps for the same two units during RUN. Centre of each row: normalized spike rate cross-correlograms (SRCs) for each state (a value of 1 represents chance, based on a uniform distribution). Note that SWS SRCs are plotted on a 20-fold narrower time scale than RUN and REM. Right of each row: SRCs calculated using the GLM method. *β* (y-axis) is the GLM coefficient for the coupling between the spike rates of the two cells. Note that SWS SRCs are plotted on a different x- and y-scale. **b:** Colour-mapped SRCs for all intramodular grid/grid cell pairs, ranked by their spatial phase offset *ϕ*_*G*_. Each row of the matrix is the colour-coded SRC of one cell pair, with red for positive cross-correlations and blue for negative cross-correlations. **b1** shows SRCs calculated using the Pearson correlation coefficient, while **b2** shows SRCs calculated according to the GLM method. The black line to the right of the matrix indicates the value of *ϕ*_*G*_ for each pair. Note the larger magnitude of negative correlations resulting from the GLM analysis. **c:** Relationship between the grid phase offset *ϕ*_*G*_ of grid/grid cell pairs and their *β*-value at zero-lag (*β*_0_). Displayed statistics are for Spearman rank (*n* = 135 grid/grid pairs)

To determine how spiking correlations varied as a function of grid phase, we sorted the GLM SRCs by *ϕ*_*G*_ and compared the ranked SRC matrices across states (**Fig. 1b2**), which revealed a consistent negative relationship between *ϕ*_*G*_ and the value of *β* at zero-lag (*β*_0_) during all states (Spearman rank *r*-values RUN -0.79, SWS -0.68, REM -0.67; all *P*-values < 10^−9^, *n* = 135 grid/grid pairs; **Fig. 1c**). We obtained a similar relationship between the zero-lag Pearson spike rate correlations and *ϕ*_*G*_(Spearman rank *r*-values RUN -0.77, SWS -0.65, REM -0.65; all *P*-values < 10^−9^, *n* = 135 grid/grid pairs; **Fig. 1b1, Supplementary Fig. 3d**), although there was an absence of strong negative correlations, in accordance with the positive bias expected from population coupling effects (**Supplementary Fig. 2a–c**). It therefore appears that the spatial tuning relationships of grid cells persist across arousal states in the form of temporal spiking correlations.

We also observed that the grid phase offset value that optimally split positive and negative correlations (*β*_0_-values) was similar during each state (RUN 0.32, SWS 0.33, REM 0.33; **Supplementary Fig. 4g**). These values closely matched the expected value determined by simulating the correlation of grid patterns as a function of phase offset (*P* > 0.05 in all cases, bootstrap, *n* = 135 grid/grid pairs; **Supplementary Fig. 4g**).

### Functional independence of modules

In contrast to the invariant spatial relationships between the grid cells within the same module, the grid patterns of cells from different modules vary independently in scale and orientation^23^. In terms of CAN models, grid modules may exist as distinct attractor networks. In such networks, activity of grid cells drawn from separate modules should be only weakly correlated across states, in comparison to correlations between cells from the same module.

We tested this prediction by identifying pairs of grid cells from different modules (**Fig. 2a–c**). We calculated SRCs for all intramodular and transmodular grid-cell pairs, during RUN, SWS and REM. Strong positive or negative peaks were generally absent from SRCs of individual transmodular cell pairs (**Fig. 2d,e**). When all GLM SRCs were ranked by the *β*_0_-value during RUN, there was a marked absence of large peaks or troughs among the transmodular SRCs compared with the intramodular SRCs, across all states (**Fig. 2f**). In confirmation of this observation, the distribution of *β*_0_ showed a significantly wider spread among the intramodular pair SRCs versus the transmodular SRCs (two-tailed Ansari-Bradley dispersion test, *P* < 0.005 during all states, *W\** < -3 during all states, *n* = 135 intramodular grid/grid pairs, *n* = 63 transmodular grid/grid pairs; **Fig. 2g**). This result was not dependent on the particular threshold chosen for classifying module membership, since we observed the same outcome across a wide range of thresholds (**Supplementary Fig. 5**).

**Figure 2:**
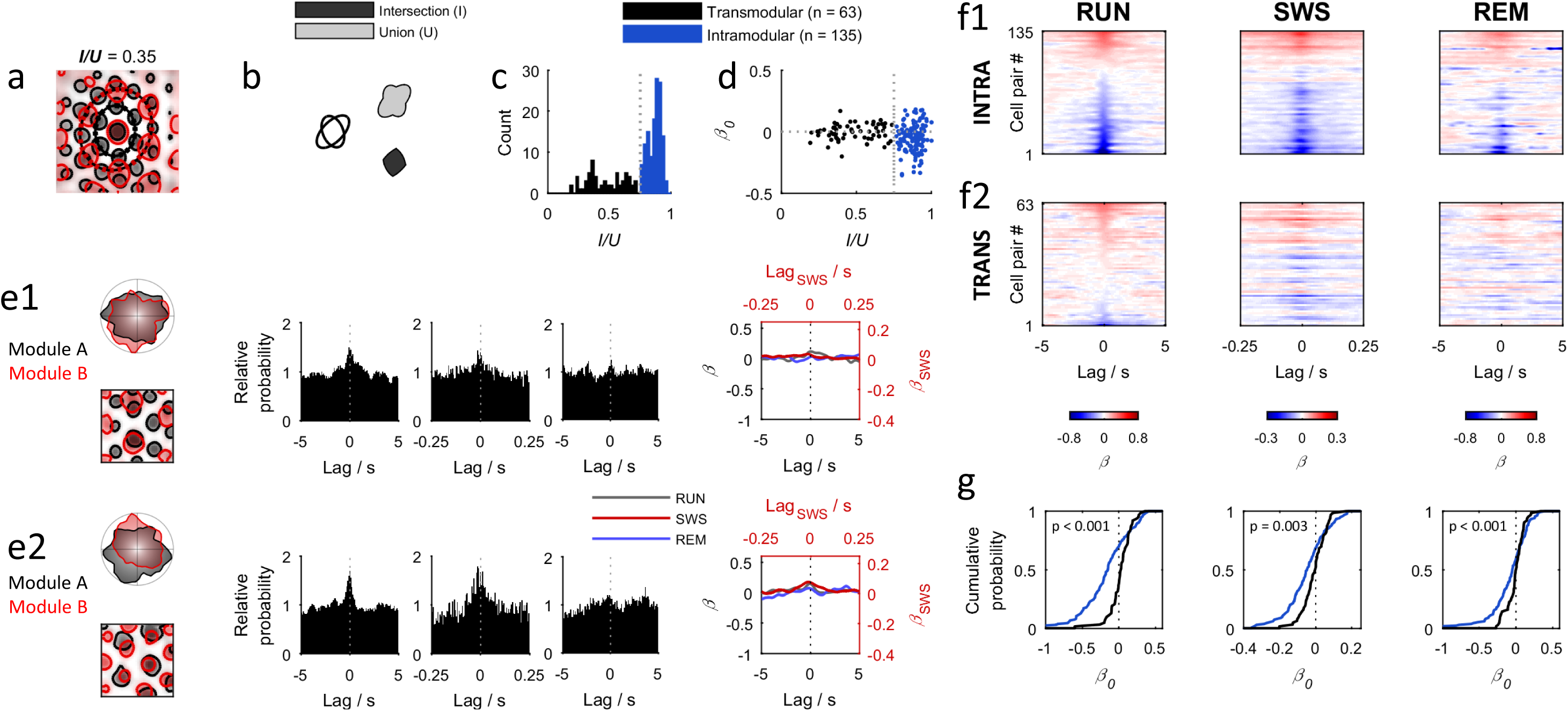
Dependence of grid-cell sleep correlations on module membership. **a**: Illustration of the module classification process. The autocorrelograms of two cells’ 2D rate maps during RUN are shown in superimposition. Ellipses are fitted to each autocorrelogram’s innermost ring of peaks, and the intersection/union area ratio (*I/U*) of the two ellipses is calculated (see (b)). The *I/U* ratio of the cell pair shown is below the threshold of 0.75 that defines them as originating from separate modules. **b**: Illustration of the intersection (dark grey) and union (pale grey) of an arbitrary pair of ellipses (black lines). **c:** Distribution of ellipse *I/U* ratios across all grid/grid pairs. The stippled vertical line indicates the module classification threshold. **d**: Relationship of grid/grid cell pairs’ GLM zero lag coupling coefficient (*β*_0_) values during SWS with the *I/U* ratio of their fitted ellipses. Note wider range of *β*_0_-values for high *I/U* ratios, in blue (intramodular comparisons). **e**: Spatial spiking patterns and spike-timing relationships of two example grid-cell pairs from different modules (**e1** and **e2**, respectively). Top left of each row: head-direction tuning curve of both units of the cell pair during RUN, one in red shade and one in grey. Bottom left of each row: superimposed 2D firing rate maps for the same two units during RUN. Centre of each row: normalized spike rate cross-correlograms (SRCs) for each state (a value of 1 represents chance, based on a uniform distribution). Note that SWS SRCs are plotted on a 20-fold narrower time scale than RUN and REM. Right of each row: SRCs calculated using the GLM method. *β* (y-axis) is the GLM coefficient for the coupling between the spike rates of the two cells. Note that SWS SRCs are plotted on a different x- and y- scale. **f:** SRCs of all intramodular (f1) and transmodular (f2) grid-cell pairs, ranked by their *β*_0_-value during RUN. Each row of the matrix is the colour-coded SRC of one cell pair, with red for positive cross-correlations and blue for negative cross-correlations. Note the wider range of values near zero-lag among intramodule pairs during all states, and the closer correspondence of the SRC values across states in the intramodular group. **g:** Cumulative distributions of *β*_0_ for intramodular and transmodular groups. Note the wider spread of *β*_0_ around zero for intramodular cell pairs in all cases. Displayed *P*-values are for two-tailed Ansari-Bradley dispersion test (*n* = 135 intramodular pairs, 63 transmodular pairs).

In summary, we find weaker correlations between transmodular grid cell pairs, consistent with the hypothesis that separate grid modules function as distinct attractor systems.

### Preservation of conjunctive grid-HD representations during sleep

The head-direction signal is a necessary precursor to the grid signal in a number of CAN models^8,18,19,33,34^. A recent study demonstrated the integrity of the network HD representation across arousal states^29^, raising the possibility that a CAN mechanism depending on HD input could maintain a similar mode of activity during offline periods.

Grid-HD cells, which conjunctively express head-directional and grid spatial tuning patterns, are a distinct class of cell that represent an interface of the HD and grid signals^8,35^. As such, they provide an opportunity to investigate the correspondence between these two signals across different brain states. If grid-HD cells inherit their HD tuning via inputs from pure HD cells, it follows that, during sleep, their spiking could remain tightly correlated with coordinated activity of the HD system^29^.

We therefore investigated how the directional tuning relationships between grid-HD and pure HD cells during RUN related to their spiking correlations across arousal states. We defined the phase difference of two HD tuning curves, *ϕ*_*HD*_, as the absolute angle between the two tuning curves’ centres of mass. Individual HD/grid-HD and HD/HD cell pairs exhibited characteristic SRC shapes, consisting of a single positive or negative peak near zero-lag whose valence was preserved across states (**Fig. 3a**). When all cell-pair SRCs were ranked by *ϕ*_*HD*_, a state-independent negative relationship between *ϕ*_*HD*_ and *β*_0_ was evident, both for HD/HD pairs^29^ (*r* < -0.7, *P* < 0.001 during all states, Spearman rank, *n =* 141 HD/HD pairs for RUN and SWS, *n* = 138 HD/HD pairs for REM; **Fig. 3b1,c1**), as previously reported, and for HD/grid-HD pairs, not reported before (*r* < -0.7 *P* < 0.001 during all states, Spearman rank, *n* = 91 HD/grid-HD pairs; **Fig. 3b2,c2**).

**Figure 3:**
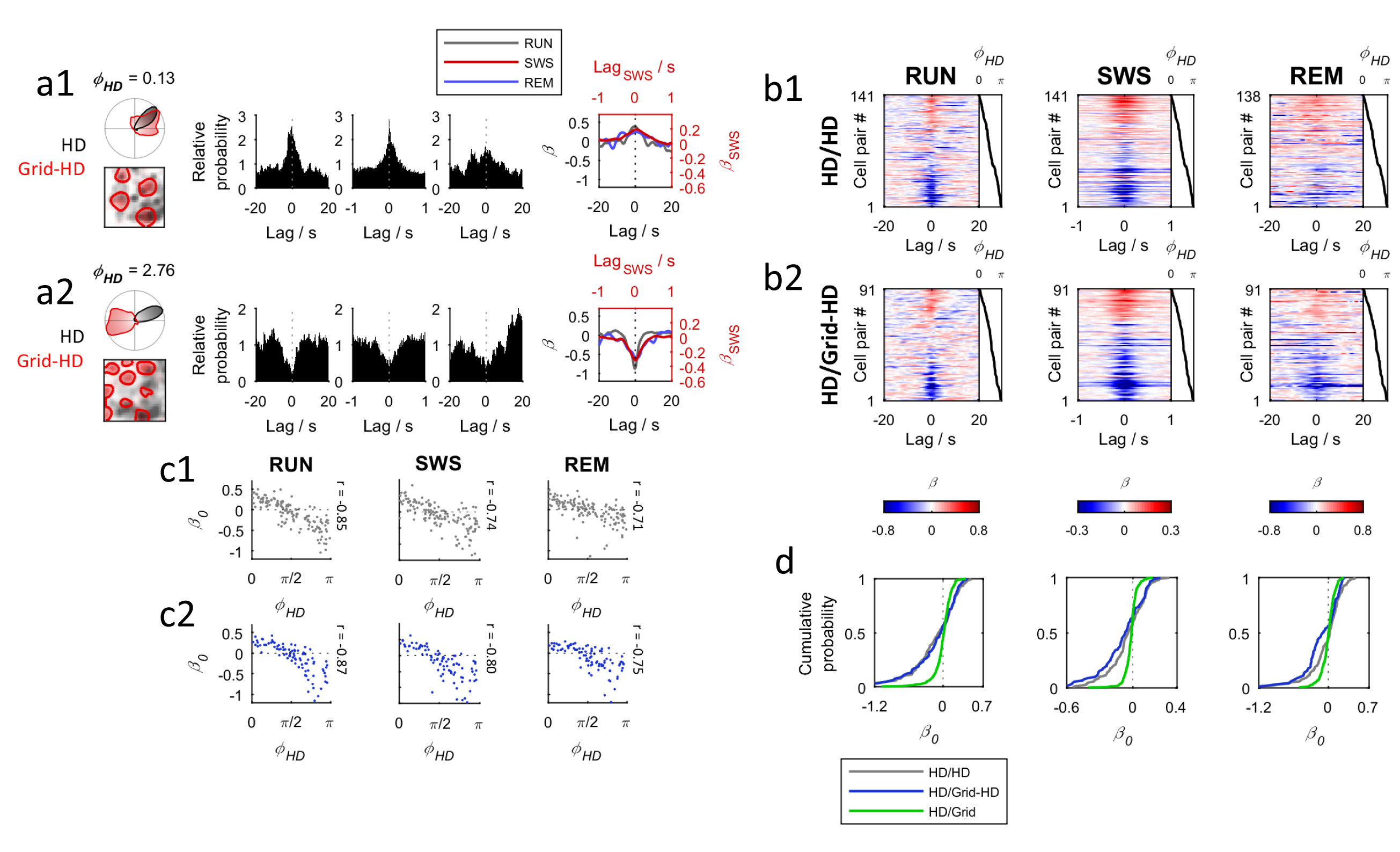
Preservation of head-directional phase offsets of conjunctive grid–HD cells during sleep. **a:** Spatial spiking patterns and spike-timing relationships of two representative HD/grid-HD cell pairs (**a1** and **a2**, respectively). Top left of each row: head-direction tuning curve of both units of the cell pair during RUN, one in red shade and one in grey. Bottom left of each row: superimposed 2D firing rate maps for the same two units during RUN. Centre of each row: normalized spike rate cross-correlograms (SRCs) for each state (a value of 1 represents chance, based on a uniform distribution). Note that SWS SRCs are plotted on a 20-fold narrower time scale than RUN and REM. Right of each row: SRCs calculated using the GLM method. *β* (y-axis) is the GLM coefficient for the coupling between the spike rates of the two cells. Note that SWS SRCs are plotted on a different x- and y- scale. **b:** Colour-mapped GLM SRCs of all cell pairs, ranked by HD phase offset *ϕ*_*HD*_. Each row of the matrix is the colour-coded SRC of one cell pair, with red for positive cross-correlations and blue for negative cross-correlations. The black line to the right of the matrix indicates the value of *ϕ*_*HD*_ for each pair. **c:** Relationship between HD phase *ϕ*_*HD*_ and GLM zero-lag coupling (*β*_0_) for HD/HD (grey) and HD/grid-HD (blue) cell pairs. Displayed *r*-values are for Spearman rank. **d:** Cumulative distributions of GLM *β*_0_-values. Across all states, the spread of *β*_0_-values is markedly wider for HD/HD or HD/grid-HD pairs, than for HD/grid pairs.

For comparison, we also computed SRCs for grid/HD cell pairs. Since these two cell types are tuned with respect to independent domains of physical space (head angle and 2D location, respectively), correlated spiking between these two cell types should not be expected during any arousal state. Accordingly, we noted an absence of strong zero-lag correlations between these two cell types during all states: the cumulative distribution of *β*_0_ was more narrowly concentrated around zero for grid/HD pairs compared with HD/HD and HD/grid-HD pairs (*P* < 10^−14^ during all states, W* < -7 during all states, two-tailed Ansari-Bradley dispersion test, *n* = 360 grid/HD pairs, *n* = 138 HD/HD pairs, *n* = 91 grid-HD/HD pairs; **Fig. 3d**). We therefore concluded that HD and grid-HD cells' directional tuning relationships are strongly preserved across arousal states, while the activity of grid cells is uncorrelated with HD cells across states.

Next, we examined the positional component of grid-HD cell tuning. Grid-HD cells coexist in modules with pure grid cells^23^; therefore we hypothesized that a fixed correlation structure should also exist between these two cell classes. Hence, we investigated the spiking correlations of intramodular grid/grid-HD cell pairs (*n* = 16; **Fig. 1a3, Supplementary Fig. 6**). The small number of cell pairs precluded conclusive analysis, and although we did observe a negative relationship between grid phase *ϕ*_*G*_ and zero-lag spiking correlation *β*_0_ during all states, this failed to reach significance during SWS (Spearman rank, RUN *r* = -0.61, *P =* 0.013; SWS *r* = -0.40; *P =* 0.129; REM *r* = -0.60; *P =* 0.015, *n* = 16 grid/grid-HD pairs). We considered that grid/grid-HD correlations may be poorly predicted by *ϕ*_*G*_ because of the additional influence of the HD signal on the spiking of grid-HD cells, which would generate more complex temporal spiking relationships. Therefore we instead used the *β*_0_-value from RUN as a predictor for *β*_0_ during SWS and REM (**Supplementary Fig. 6c,d**). In this case, we found a strong correlation between *β*_0_ during RUN and *β*_0_ during SWS/REM (*r* = 0.81 for both RUN-SWS and RUN-REM; *P* < 0.001, Spearman rank, *n* = 16 grid/grid-HD pairs). Thus, while *ϕ*_*G*_ is insufficient to predict grid/grid-HD correlations across states, there is nevertheless a persistence of correlation structure across states.

### Intra-state stability of correlation structure

Although we found overall grid, grid-HD and HD spiking correlations to be highly conserved across three widely different brain/behaviour states, it is not clear whether these correlations manifest at all times, or are restricted to particular time windows. To probe for short-term correlation variability, we examined how the spiking correlations between pairs of MEC cells varied over time during SWS epochs. We isolated all cell pairs that were recorded during rest sessions containing at least 60 minutes of SWS (*n* = 101 intramodular grid/grid and *n* = 102 HD/HD). For each cell pair, we calculated SRCs from spikes within a 2-minute moving window shifted in 1-minute steps (**Fig. 4**). The shapes of time-resolved SRCs showed remarkably little departure from the aggregate SRCs.

**Figure 4:**
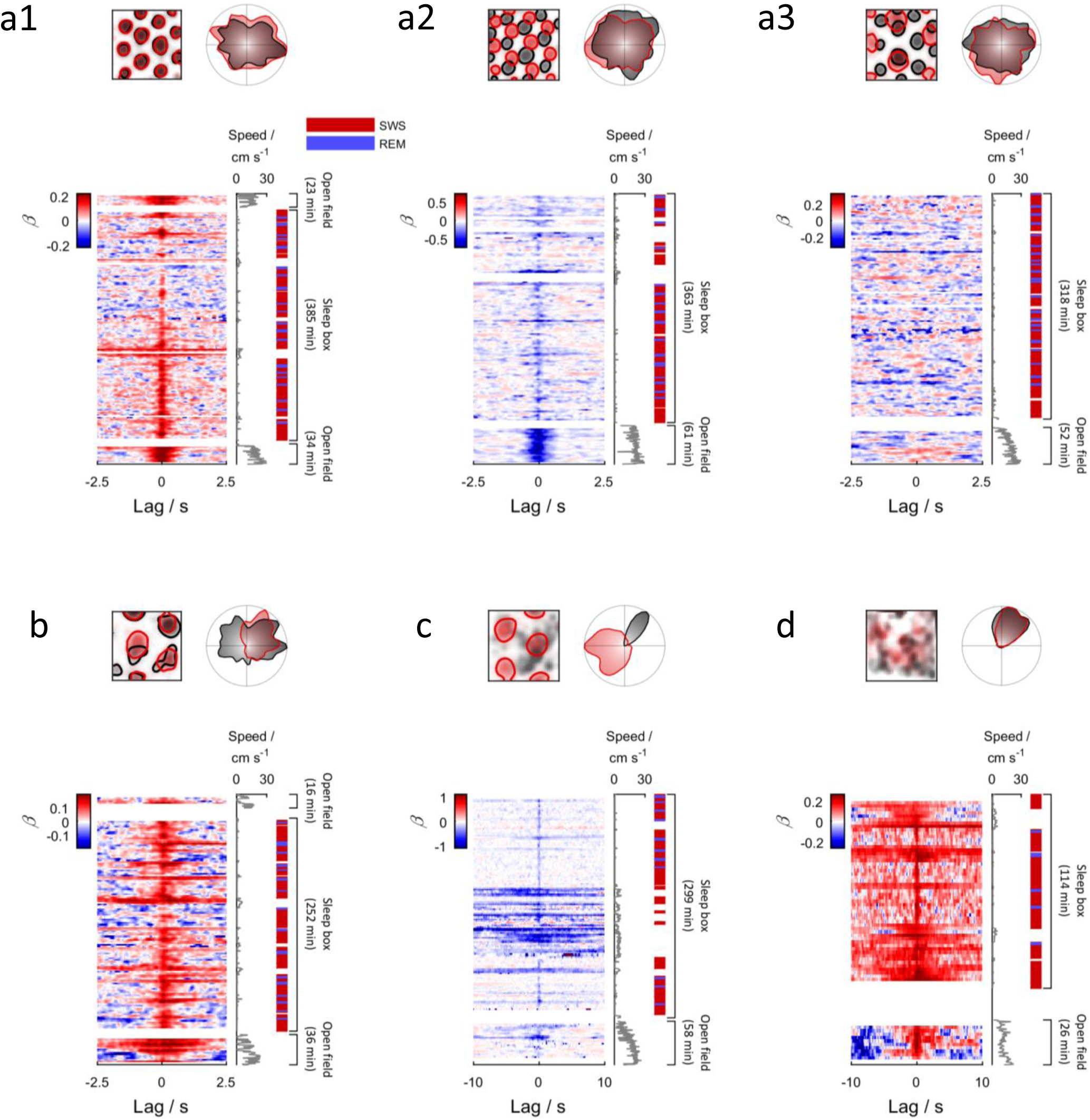
Temporal stability of correlations: example cell pairs. **a1-a3**, Example time-resolved SRC plots for 6 cell pairs (one panel each). Top of each panel: 2D firing rate maps (left) and HD tuning curves (right) calculated during RUN, with one cell in red shade and one cell in grey shade; bottom of each panel: time-resolved SRCs calculated in 2-minute bins across the whole recording. Each row of the matrix represents the SRC for one 2-minute bin, with the bottom row representing the beginning of the recording. Cross-correlation (*β*) is colour-coded. Time bins with fewer than 100 counts in the SRC histogram were excluded (blank rows). The grey line to the right of the plot indicates the movement speed of the animal. The coloured blocks to the right of the plot indicate periods of SWS (red) and REM (blue). **a** shows three pairs of pure grid cells (**a1**, in-phase; **a2**, out-of-phase; **a3**, separate modules). Note the persistence of zero-lag correlations in a1/a2, and the absence of correlation in a3. **b** shows a pure grid cell paired with a grid-HD cell; **c** shows a HD cell paired with a grid-HD cell; **d** shows two HD cells.

To quantify the variability of correlations across SWS epochs, we merged the activity from SWS epochs together and divided the merged activity into 15-minute blocks, for each of which we calculated *β*_0_ for each cell pair. To quantify coupling variability, we compared the GLM *β*_0_ vaues for the first and last 15-minute windows, using the correlation between these two variables as a measure of coupling stability. Among grid/grid and HD/HD cell pairs, the *β*_0_-values from the two time windows were strongly correlated (grid/grid pairs, *r*= 0.80, *P* < 10^−8^, Spearman rank, *n* = 100; HD/HD pairs, *r* = 0.89, *P* < 10^−8^, Spearman rank, *n* = 97; **Fig. 5a**), indicating high stability of grid and HD correlation structure across SWS.

**Figure 5:**
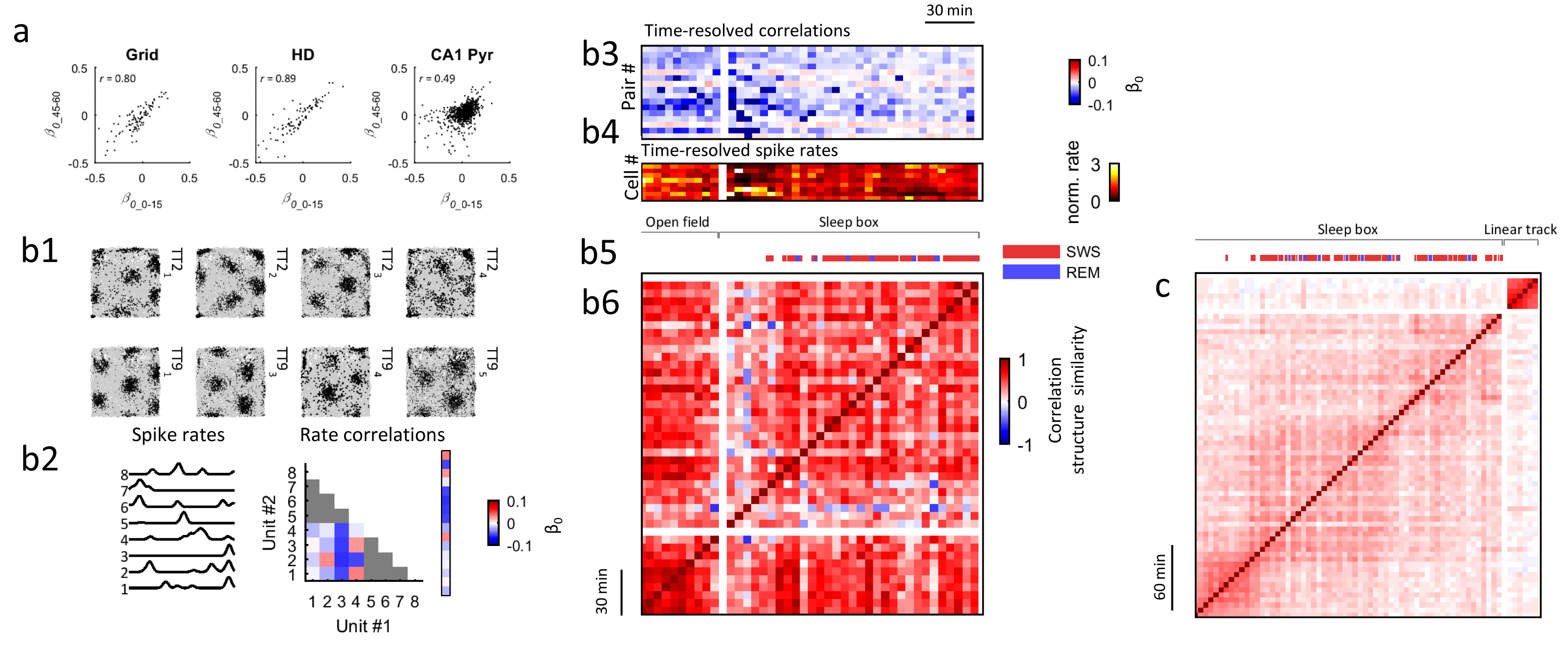
Within-state stability of correlation structure. **a:** Stability of GLM *β*_0_-values between grid/grid, HD/HD and CA1/CA1 cell pairs. *β*_0_-values were calculated separately for the first and fourth 15-minute blocks of concatenated SWS (0–15 and 45–60 minutes). These two *β*_0_-values are shown for each cell pair (dot) on the x- and y-axes respectively. Note the substantially higher stability of grid/grid and HD/HD correlations compared with CA1/CA1 correlations. Displayed *P*-values are for Spearman rank correlation (*n* = 100 grid/grid pairs, 97 HD/HD pairs, 934 CA1/CA1 pairs). **b:** Time course of correlation structure in an example ensemble of 8 grid cells from the one module. **b1:** Locations of spikes for each neuron during RUN (rat’s trajectory in grey, spike locations in black). **b2:** Calculation of pairwise zero-lag GLM *β*_0_-values. Left: spike rates of the eight cells. Centre: matrix showing *β*_0_-values for pairs of cells. Cell pairs originating from the same tetrode are not counted and are greyed out. Right: vector containing all *β*_0_-values from the correlation matrix. **b3:** Matrix composed of *β*_0_-values for all grid-cell pairs, calculated in 5-minute time bins. Cell pairs are in rows (in arbitrary order), time bins are in columns. Note persistence of *β*_0_- values over time**. b4:** Spike rates of the 8 grid cells across the same time range plotted in (b3). **b5:** Brackets marking the boundaries of different sessions within the recording, and identified sleep periods during the sleep box session (coloured bars). **b6:** Metacorrelation matrix representing pairwise correlation structure within the 8-cell ensemble. Each row or column in the matrix represents the vector of *β*_0_-values at the corresponding time bin in (b3), and each matrix element represents the Pearson *r*-value yielded by correlating the two *β*_0_ vectors corresponding to its row and column indices. Note the predominance of strongly positive values (red colour) across the entire matrix, indicating global preservation of the ensemble spiking correlation structure. **c:** Same as (b6), but generated from a sample of 50 concurrently recorded CA1 pyramidal cells. Note the much lower general preservation of correlation structure across time periods in comparison to (b6).

A major difference between entorhinal and hippocampal spatial maps is the low dimensionality of the former and the high dimensionality of the latter^36^. Unlike grid cells^22–24^, head direction cells^37–39^, and border cells^12^, hippocampal place cells remap between contexts and environments in a manner that often generates completely orthogonal maps for very similar environments, even when the number of environments compared is extensive^40–42^. This difference led us to compare the fixed activity structure of the MEC network during sleep with activity in hippocampal area CA1. Using a set of previously published recordings of CA1 putative pyramidal cells during rest before a linear track running session (see methods)^43^, we applied the same analyses as for the MEC data from the present recordings. Due to the large number of cells in some recordings, we selected a maximum of 200 cell pairs at random from each recording (*n* = 6 recordings, *n* = 934 cell pairs total). In contrast to grid/grid and HD/HD cell pairs, CA1/CA1 cell-pair couplings showed weak stability between the first and fourth 15-minute SWS blocks (*r* = 0.49, *P* < 10^−10^, Spearman rank, *n* = 934 CA1/CA1 pairs; **Fig. 5a**), indicating more variable CA1 pyramidal correlations during SWS (*P* < 0.001 compared with grid/grid and HD/HD correlations, bootstrap, *n* = 100 grid/grid pairs, *n* = 97 HD/HD pairs, *n* = 934 CA1/CA1 pairs). Indeed, there was not a single recording in which the CA1 correlation stability reached that of HD or grid cells (highest *r*-value 0.63; **Supplementary Fig. 7c**). The lower stability of CA1/CA1 pair correlations was preserved when all units’ spike rates were subsampled to 0.2 Hz, in comparison to both grid/grid pairs and HD/HD pairs (bootstrap, *n* = 934 CA1/CA1 pairs, 100 grid/grid pairs, 97 HD/HD pairs), ruling out differences in mean spike rate as a possible cause. Furthermore, the effect was robust across a wide range of rate smoothing kernel widths (**Supplementary Fig. 7b**).

To examine grid-cell correlation stability at a wider network level, we examined the ensemble activity of 8 concurrently recorded grid cells from a single module (**Fig. 5b1**). We divided the recording into nonoverlapping 5-minute time windows, and within each window created a vector comprising the set of *β*_0_-values for all cell-pair combinations within the ensemble (**Fig. 5b2–3**). From these *β*_0_-value vectors, we generated a matrix of metacorrelations by correlating the *β*_0_-value vectors at every pair of time points (**Fig. 5b6**). This metacorrelation matrix was predominated by strongly positive *r*-values, indicating a global preservation of the spiking correlation structure among the 8 grid cells, measured at the network level. We also performed this analysis on an ensemble of 50 CA1 pyramidal cells (**Fig. 5c**). The metacorrelation matrix was predominated by much smaller positive values, indicating less global stability of correlation structure.

In summary, we found that grid and HD correlations show similarly strong intra-state stability, while CA1 pyramidal correlations are comparatively dynamic.

### Effect of hippocampal sharp-wave-ripple activity on correlations

The CA1 region of hippocampus provides a major input to to the deep layers of MEC^44,42,45–47^, and synchronous population events generated in hippocampus known as sharp-wave-ripples (CA1-SWR) have been shown to shortly precede activation of deep-layer MEC neurons^48,49^. To investigate the possibility that correlated spiking between grid cells could be due to selective coactivation by CA1- SWR, we examined the relationship between pairwise spiking correlations and CA1-SWR activity.

CA1-SWRs have been shown to induce sharp-wave deflections in MEC (MEC-SW)^49,50^. We identified MEC-SW which were correspondent with CA1-SWR events (2 rats; **Fig. 6a–c**). Higher-amplitude CA1- SWR events were associated with higher-amplitude MEC-SW events (**Fig. 6c**). For each single unit, we calculated its average spike rate in a ±0.25 s window around MEC-SW. In each class of units, a large proportion of cells showed strong rate modulation by MEC-SW occurrence, with peak rates typically falling within 20 ms of the MEC-SW (latency of peak in mean perievent rate: grid +3 ms, HD -12 ms, grid-HD -7 ms; *n* = 95 HD cells, 134 grid cells, 39 grid-HD cells; **Fig. 6d,e**).

**Figure 6:**
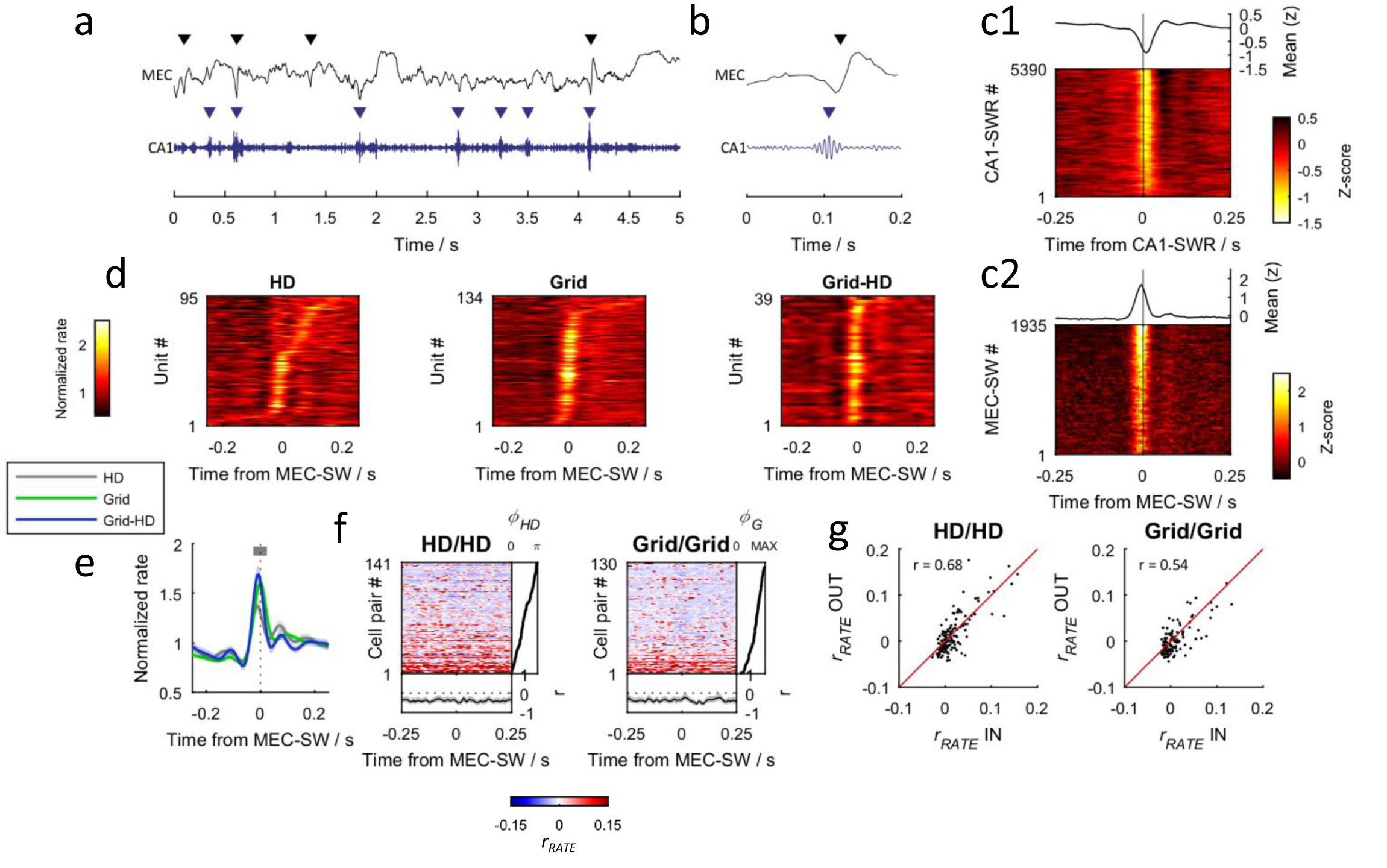
Effect of MEC sharp waves on spike rates and correlations. **a:** Example trace of MEC and CA1 LFP activity during SWS, showing wideband MEC LFP (top) and 125- 250 Hz bandpass-filtered CA1 LFP (bottom). The CA1 LFP contains brief periods of high-amplitude 125–250 Hz activity known as sharp-wave-ripples (CA1-SWR), marked with blue arrows, while the MEC LFP contains prominent sharp waves (MEC-SW), marked with black arrows. Note that two of the MEC-SW events coincide with CA1-SWR events. **b:** Example trace of MEC and CA1 LFP activity during a single CA1-SWR /MEC-SW event. **c:** Event-triggered LFP traces from an example recording. **c1** shows MEC LFP traces anchored to the times of CA1-SWR events detected during SWS. Events are ordered from the lowest amplitude (bottom) to the highest amplitude (top). The average MEC LFP response is shown above. **c2** shows CA1 LFP amplitude within the 125–250 Hz band, anchored to the times of identified MEC-SW events detected during SWS. Events are ordered from the lowest amplitude (bottom) to the highest amplitude (top). The average 125–250 Hz amplitude response is shown above. **d:** Time course of spike rates anchored to MEC-SW events. Each row represents the mean MEC-SW-triggered spike rate of one unit. Firing rate is colour-coded. Rates were normalized by dividing by the mean value across the perievent time window. Units are displayed ordered by the timing of their maximal firing rate. **e:** Mean MEC-SW-triggered spike rates across all units (*n* = 95 HD cells, 134 grid cells, 39 grid-HD cells). The grey bar indicates the time range defining the MEC-SW period. Shaded areas represetn 95% bootstrap confidence intervals. **f:** Time course of pairwise spike rate Pearson correlations (*r*_*RATE*_) anchored to MEC-SW events, for HD/HD (left) and grid/grid pairs (right). Pairs are ordered by their phase offset, from smallest (bottom) to largest (top). The black line to the right of the matrix indicates the phase offset of each cell pair. The black line beneath the matrix indicates the Spearman rank correlation between *r*_*RATE*_ at each time step and the cell-pair phase offset values (*n* = 141 HD/HD pairs, 130 grid/grid pairs; the shaded area shows 95% bootstrap confidence intervals). Note the lack of perturbation of *r*_*RATE*_-values at zero-lag. **g:** Comparison of *r*_*RATE*_ during MEC-SW epochs (IN), versus *r*_*RATE*_ outside of MEC-SW epochs (OUT). The IN period corresponds to the range shown by the grey horizontal bar in (e); the OUT period corresponds to the remaining part of the ±0.25 s perievent window. Displayed *r*-values are Spearman rank (*n* = 141 HD/HD pairs, 130 grid/grid pairs).

Having observed that all cell types showed substantial spike rate modulation by MEC-SW, we went on to examine how MEC-SW impacted pairwise spiking correlations. For each HD/HD and grid/grid cell pair, we calculated spike rate correlations iteratively across a ±0.25 s window around MEC-SW (**Fig. 6f**). Due to the limited number of MEC-SWs available, which resulted in unreliable fitting for the GLM, we reverted to using the conventional Pearson correlation coefficient as the correlation metric (*r*_*RATE*_). To quantify the impact of MEC-SW on pairwise spike rate correlations, we defined a ±25 ms window around the sharp-wave peak which encompassed the times when all cell types reached their peak mean rates (**Fig. 6e**). For both HD/HD and grid/grid cell pairs, *r*_*RATE*_ appeared unperturbed by MEC-SW (**Fig. 6f**), and was similar between MEC-SW and non-MEC-SW periods (HD/HD pairs *r* = 0.68, Spearman rank, *n* = 141; grid/grid pairs *r* = 0.54, Spearman rank, n = 130; **Fig. 6f,g**), despite the large effects on single unit spike rates (**Fig. 6d,e**). For both grid/grid and HD/HD pairs, spatial phase offsets were similarly correlated with spike rate correlations during (IN) and outside (OUT) MEC-SW periods (HD/HD pairs IN: *r* = -0.580, *P* < 10^−10^, OUT: *r* = -0.676, *P* < 10^−10^, Spearman rank, *n* = 141; grid/grid pairs IN: *r* = -0.390, *P* < 10^−5^, OUT: *r* = -0.594, *P* < 10^−10^, Spearman rank, *n =* 130; no difference between IN/OUT *r*-values for both HD/HD and grid/grid pairs, *P* > 0.05, bootstrap, *n* = 141 HD/HD pairs and 130 grid/grid pairs; **Fig. 6f**). In conclusion, the HD- and grid-cell correlations described in the present study appear to persist irrespective of the occurrence of MEC-SWRs.

### Temporal dynamics of latent states

Spike rate cross-correlations contain information about the temporal dynamics of the signals that influence the cells’ spiking^29,51,52^. Since our observations of preserved correlation structure are consistent with grid module networks containing a covert grid representation during sleep, we reasoned that we should be able to infer the dynamics of such a signal by using a generative model to reconstruct spiking cross-correlations^29^. Such a reconstruction would show whether a covert grid pattern would generate a structure of pairwise cross-correlations similar to the one observed in our sleep data.

Adapting an approach previously applied to HD cells^29^, we modelled the covert grid signal as a 2- dimensional random-walk process (see methods; **Fig. 7a**). For each pair of intramodular grid cells, we simulated the cells' spike rates in response to the random walk, and generated SRCs from the simulated spike rates (**Fig. 7b**). We iteratively simulated SRCs using a wide range of random walk drift rates, and determined which speed yielded the closest fit to the original SRCs from both RUN and SWS. The best-fit model SRCs closely approximated the shape of the original SRCs (**Fig. 7b,c**). The median best-fit drift rate markedly increased from 1.69 cycles s^-1^ (where one cycle corresponds to the grid spacing) during RUN to 8.40 cycles s^-1^ during SWS (*P* < 10^−13^, sign-rank test, *n* = 85 grid/grid pairs), with the best-fit drift rates during SWS being a median of 4.98 times higher than during RUN (**Fig. 7g,h**).

**Figure 7:**
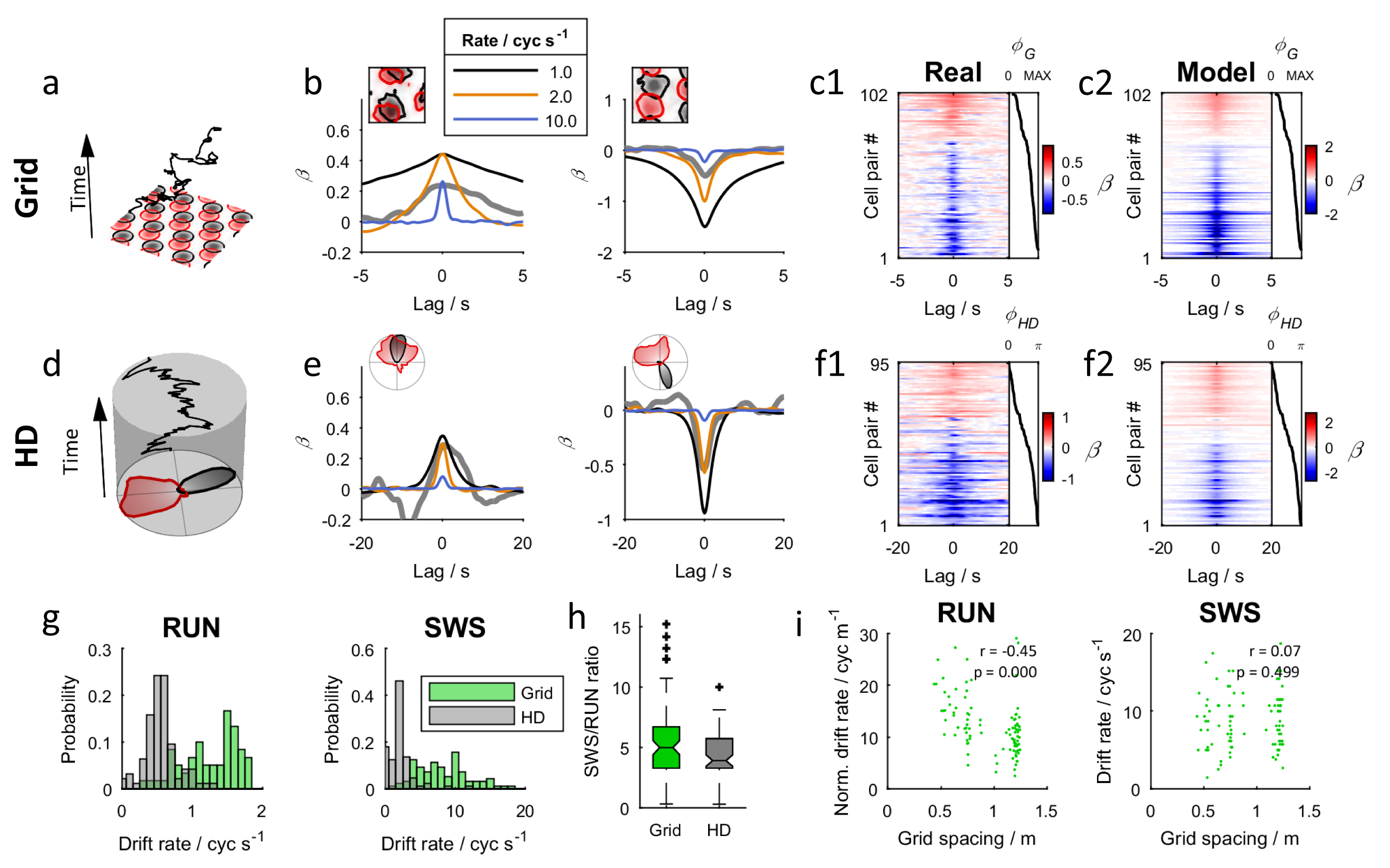
Comparison of temporal dynamics in grid and HD signals in RUN and SWS. **a:** Illustration of a two-dimensional random walk through 2D space (black line). Time is represented as increasing along the vertical axis. The red and black grid patterns indicate the receptive field locations of two simulated grid cells. **b:** SRCs for two grid/grid pairs calculated during SWS. The rate maps for the grid/grid cell pair are shown in the inset (left pair in-phase, right pair out-of-phase). In the line plot below, the thick grey line shows SRCs for recorded data in RUN, whereas the coloured lines indicate simulated SRCs generated using random walks with mean drift rates of 1, 2 or 10 cycles per second. One cycle corresponds to the grid spacing. **c:** Colour-mapped GLM SRCs of all cell pairs, ranked by grid phase offset *ϕ*_*G*_. Each matrix row is the SRC of one cell pair. The black line to the right of the matrix indicates the value of *ϕ*_*G*_ for each pair. Cross-correlation (*β*) is colour-coded. **c1** shows the original SRCs; **c2** shows the best-fit reconstruction from the random-walk model. Note that the original SRCs generally have *β*-values smaller in magnitude than the random-walk reconstructions. This effect is likely due to population rate coupling in the original spike trains, which is not incorporated into the random-walk model. **d:** Illustration of a random walk in the head-direction domain. The tuning curves of an example pair of HD cells are shown. The angles of the random walk are plotted on a cylinder, with time as the vertical axis. **e:** Same as (b), but for two pairs of HD cells (one pair in-phase, one out-of-phase). One cycle corresponds to a full 360-degree rotation. **f:** Same as (c), but for HD/HD cell pairs, ranked by HD phase offset *ϕ*_*HD*_. **g:** Distribution of estimated drift rates for grid/grid and HD/HD pairs during RUN and SWS. Note the faster cycling of grid cells during both states. **h:** Box plot for estimated SWS/RUN drift rate ratio. Box indicates median and quartiles; whiskers indicate 1.5× interquartile range (*n* = 85 grid/grid pairs, 66 HD/HD pairs). **i:** Relationship between grid spacing and estimated drift rate, during RUN and SWS. For RUN, variations in the running speed of the animal are accounted for by dividing each drift rate by the animal’s mean running speed during the RUN session (“norm. drift rate”). For SWS, the raw drift rate is shown. Drift rates are strongly correlated with grid spacing during RUN, but become uncoupled during SWS. The displayed statistics are for Spearman rank correlation (*n* = 100 grid/grid pairs during RUN, 102 grid/grid pairs during SWS).

During wakeful navigation, each grid module represents an animal's location in its environment using a grid pattern with a single spacing^23^. When the animal moves at a particular speed, grid signals encoded in larger-scale modules must show a slower temporal rate of change than grid signals encoded by smaller-scale modules; thus there is a direct relationship between spatial and temporal scales of grid cell activity. It is not clear whether a scale relationship exists between grid modules during sleep, so we addressed this question by comparing the grid spacing of grid/grid cell pairs with their best-fit random walk drift rate. During RUN, as expected, drift rate was negatively correlated with grid spacing (*r* = -0.449, *P* = 2.19 x 10^−6^, Spearman rank, *n* = 102 grid/grid pairs; **Fig. 7i**). In contrast, we observed no such relationship during SWS (*r* = 0.0684, *P =* 0.499, Spearman rank, *n* = 100 grid/grid pairs), suggesting that there is no temporal scale relationship between modules during SWS.

So that we could compare the temporal dynamics of the grid and HD representations during SWS, we also applied the above analysis to pairs of HD cells by modelling the covert HD signal as a 1- dimensional circular random walk process (**Fig. 7d**). As with grid cell pairs, we found that random walks yielded good fits to the observed HD/HD SRCs (**Fig. 7e,f**). During both RUN and SWS, the estimated drift rates of HD/HD cell pairs were substantially lower than grid/grid pairs (median drift rates RUN 0.599 cycles s^-1^, SWS 2.42 cycles s^-1^, where 1 cycle corresponds to a full 360° rotation; comparisons with grid drift rates *P* < 10^−26^, rank-sum test, *n* = 85 grid/grid pairs, 66 HD/HD pairs, z > 10). The median SWS:RUN acceleration factor of 3.90 was similar to that of grid/grid cell pairs (*P =* 0.107, rank-sum test, *n* = 85 grid/grid pairs, *n* = 66 HD/HD pairs, z = 1.61; **Fig. 7h**), indicating a parallel change in temporal dynamics of the grid and HD systems between alert wakefulness and SWS.

## Discussion

We have shown that spatial tuning relationships between grid cells are upheld across brain states in the form of a rigid temporal spiking correlation structure. The preserved network states extended to conjunctive grid-HD cells, suggesting that the functional relationship between the HD and grid networks is also fixed. The highly constrained activity that these correlations signify is a signature of continuous attractor network dynamics, and therefore lends support to hypotheses of grid formation based on such mechanisms.

### Persistent constraints on grid cell activity

We demonstrated that the spatial tuning relationships between pairs of grid cells during open-field navigation are accompanied by spiking correlations which remain fixed between arousal states. Furthermore, the pairwise correlations between grid cells varied little within SWS, in stark contrast to CA1 pyramidal cell correlations, which were highly dynamic. The correlations in MEC were notably strong, considering the short timescale on which they were measured and the low mean spike rate of the cells^53^, which points to forceful constraints on coordinated activity. Although our observations were principally based on pairwise correlations rather than large-scale ensemble activity, it is difficult to conceive how a strong pairwise correlation structure could exist in the absence of coordination at the population level. Therefore, the most parsimonious interpretation of the persistent pairwise correlation structure between grid, grid-HD and HD cells is that these networks behave according to low-dimensional intrinsic manifolds that result from selective, fixed synaptic connectivity.

Previous attempts to examine correlation structure in the grid-cell network have relied on analyses that subtracted inferred signal components from correlations in order to measure the residual noise correlations^32,54^. In these studies, particular assumptions had to be made about the nature of signal correlations, and therefore it is possible that signal correlations were not fully accounted for. We circumvented the issue of separating signal and noise correlations by focussing on a state during which spatial signals are absent. Our observations are reminiscent of the recent discovery that HD network activity remains constrained on a 1-dimensional circular manifold across all states^29^. In that study as well as our own, cells’ spatial tuning relationships were tightly linked to invariant temporal spiking correlations across arousal states. It was demonstrated in the previous study that during SWS and REM, HD network activity coherently represented a covert HD signal that drifted continuously and independently of the animal’s actual head direction. It is plausible that a similar covert grid signal exists during sleep. Due to the additional dimension represented in the grid signal, precise decoding requires many more cells than is required to decode the HD signal, and more than was available in the present work. Yet we demonstrated through modelling that random-walk dynamics of a covert grid signal were sufficient to replicate grid cells' spiking cross-correlation patterns during wake and SWS.

Conjunctive grid-HD cells, which are abundant in layer 3 of MEC^35^, provide a valuable opportunity to determine the relationship between HD and grid representations. Remarkably, we found that the conjunctive nature of these cells was maintained during sleep: they demonstrated spiking correlations with pure HD cells as prescribed by their HD phase offset, while the shape of their spiking correlations with grid cells was also maintained across states. As such, our findings are consistent with conjunctive grid-HD cells functioning as an interface between covert HD and grid signals during sleep, in a fashion which would parallel their activity during navigational behaviour^8^.

During SWS, we observed a 4–5-fold acceleration of grid and HD correlation temporal dynamics. This is consistent with prior accounts of activity in other forebrain regions, which generally point to an acceleration of neural dynamics during SWS ^29,51,52,55^. REM correlations, conversely, showed a temporal scale that was similar to RUN, reinforcing the notion that REM mimics the general dynamics of wakefulness in the entorhinal-hippocampal and HD systems ^29,56,57^.

### Relationship to continuous attractor network models

A number of theoretical models for grid pattern formation are based on continuous attractor network mechanisms ^8,18,19,33,58^. A universal feature of CAN models is that selective recurrent synaptic connectivity forms the basis for the spatial phase offsets between grid cells: grid cells with the strongest net excitatory connectivity have the smallest spatial phase offsets. Therefore, an important prediction of CAN models is that functional connectivity should exist between grid cells as a function of their spatial phase offset, irrespective of the network’s mode of activity. In support of these predictions, previous work has demonstrated fixed spatial relationships between the receptive fields of grid cells within a module ^22–24^. The present study adds to those observations by revealing that constraints on grid cell spiking relationships span not only different spatial environments, but also different brain states. This observation is difficult to explain in terms other than a fixed synaptic basis for grid-cell phase offsets, which underlies all CAN models of grid cells. Our study also showed that grid modules act independently during sleep, with similar temporal dynamics irrespective of the spatial scale of the modules. This is in accordance with the hypothesis that grid modules exist as distinct attractor networks, a necessary requirement of all CAN models for grid cells^8,20^.

The mechanisms of CAN models vary widely, however, and a natural further question is whether our present findings specifically lend support to any subset of CAN models. One salient aspect of our findings is the relationship between the HD and grid signals. In some CAN grid models, the HD signal is a crucial input which, together with a signal encoding the animal’s speed ^59,60^, is used to translate the grid pattern in correspondence with the animal’s movement ^8,18,19,33,58^. Conjunctive grid-HD representations may emerge as intermediate steps in the computation of the “pure” grid signal^8^. Our findings suggest that an interface between the HD and grid signals remains during sleep, since grid-HD cells retained their correlations with both pure grid and HD cells. This would be consistent with a path-integrating CAN model in which the HD signal is an essential precursor. It should be added, however, that our observations do not directly rule out alternative mechanisms, such as feedforward Hebbian self-organization mechanisms^61^, as long as the fixed activity structure of the MEC networks is formed prior to experience, during early development of the nervous system^62^. However, in their native form, these alternative models predict neither the independent operation of the grid modules nor the coherence of grid cells and grid-HD cells.

### Functional implications

While the present results are predicted by CAN grid models, the functional role that a grid-like network representation might serve during sleep, if any, is not clear. One possible mode of activity of the offline grid signal could be to “replay” patterns of neuronal activity that were previously established during spatial navigation^63^. Replay has been studied in detail in the hippocampus, where, during restful periods, place cells reactivate in brief and rapid sequences that follow the same temporal order that they were activated in a particular locomotive trajectory ^51,55,64,65^. More recently, it has been reported that hippocampal replay is accompanied by synchronized reactivations of homologous trajectories in grid cells in deep layers of MEC ^48^. It has additionally been claimed that superficial MEC cell ensembles, including grid and non-grid cells, can generate sequences that represent navigational trajectories, independently of hippocampal activity ^30^. However, whether sequence replay takes place in MEC ensembles remains to be determined, as concerns have been raised over the methodological complications of detecting replay in grid cell populations ^66,67^. Furthermore, we observed a loss of module scale relationships during SWS, which is incompatible with coordinated replay across modules.

In the present study, we do not directly address the phenomenon of replay but the findings have implications for the interpretation of replay-like activity in MEC. Our study shows that continuous attractor dynamics provide a good description of coordinated grid-cell activity across different arousal states. The constrained nature of the grid-cell network contrasts starkly with the hippocampal place cell network, whose correlation structure is highly dynamic by comparison, undergoing orthogonalization across different contexts ^40–42,68^, and showing comparatively low stablility of spiking correlations during SWS in the present results. This fundamental difference between the grid- and place-cell networks has important methodological implications for detecting replay. Assessing the significance of sequential spiking events typically involves randomly shuffling neuronal activity to generate a null distribution of sequences that would be expected to occur by chance ^56,65,69^. In the hippocampus, shuffling methods that independently permute activity between place cells may generate a valid null distribution, since the randomized sequences may all be equally permissible during stochastic behaviour of the hippocampal network. However, in the grid cell network, the same shuffling methods would overwhelmingly generate “off-manifold” population states that would violate the network's low-dimensional constraints, and therefore would not constitute a valid null hypothesis. Because employing such shuffling methods would inevitably favour the detection of false-positive replay events, we cannot exclude that prior observations of grid-cell replay that relied on cell-independent shuffling methods reflect purely stochastic network processes. It will be vital for future studies of replay in MEC to employ statistical approaches that respect the low-dimensional constraints on grid-cell population activity.

## Methods

### Subjects

Data were collected from 7 male Long Evans rats, which were experimentally naive and 3–5 months old (350–600 g) at the time of implantation. The rats were group-housed with 3–8 of their male littermates prior to surgery, and were singly housed in large Plexiglas cages (45 x 44 x 30 cm) thereafter. The rats were kept on a 12 hr light/12 hr dark schedule, and humidity and temperature were strictly controlled. The experiments were performed in accordance with the Norwegian Animal Welfare Act and the European Convention for the Protection of Vertebrate Animals used for Experimental and Other Scientific Purposes.

### Electrode Implantation and Surgery

Tetrodes were constructed from four twisted 17 μm polyimide-coated platinum-iridium (90%/10%) wires (California Fine Wire). The electrode tips were plated with platinum to reduce electrode impedances to between 120–300 kω at 1 kHz.

Anaesthesia was induced by placing the animal in a closed plexiglas box filled with 5% isoflurane vapour. Subsequently, the animal received a subcutaneous injection of buprenorphine (0.03 mg kg^-1^), atropine (0.05 mg kg^-1^) and meloxicam (1.0 mg kg^-1^), and was mounted on a stereotactic frame. The animal’s body rested on a heat blanket to maintain its core body temperature during the surgical procedure. Anaesthesia was maintained with isoflurane, with air flow at 1.0 litres/minute and isoflurane concentration 0.75–3% as determined according to breathing patterns and reflex responses.

The scalp midline was subcutaneously injected with the local anaesthetic lidocaine (0.5%) prior to incision. After removal of the periost, holes were drilled vertically in the skull, into which screws (M1.4) were inserted. Two screws positioned over the cerebellum were used as the electrical ground. Craniotomies were drilled anterior to the transverse sinus. Subsequently, the animal was implanted with either a hyperdrive containing 14 independently moveable tetrodes (six animals), or two microdrives, each containing a single bundle of 4 tetrodes (one per hemisphere; one animal). Hyperdrive implants were always on the left side. Hyperdrive tetrodes were implanted between 3.75 and 5.0 mm ML, with the posterior edge of the bundle located 0.1–0.2 mm anterior to the edge of the transverse sinus. Each tetrode was immediately advanced by 960–1280 μm. Microdrive tetrodes were inserted at 4.5 mm ML, 0.1–0.15 mm anterior to the transverse sinus edge at an angle of 20 degrees from vertical in the saggital plane. The tetrodes were immediately inserted to a depth of 1800 μm. Two animals with hyperdrive implants were also implanted with a single-wire LFP electrode in hippocampus CA1 (-4.0 mm AP, 3.2 mm ML, 2.2–2.4 mm ventral to brain surface). Implants were secured with dental cement (Meliodent). 8–12 hours after the beginning of the surgery, the animal was treated with an additional dose of buprenorphine (0.03 mg kg^-1^).

### Recording procedures

Over the course of 1–3 weeks, tetrodes were lowered in steps of 320 μm or less, until high-amplitude theta-modulated activity appeared in the local field potential at a depth of approximately 2.0 mm. In hyperdrive experiments, two of the tetrodes were used to record a reference signal from white matter areas. The drive was connected to a multichannel, impedance matching, unity gain headstage. The output of the headstage was conducted via a lightweight multiwire tether cable and through a slip-ring commutator to a Neuralynx data acquisition system (Neuralynx, Tucson, AZ; Neuralynx Digital Lynx SX, for all hyperdrive-implanted animals), or via a counterbalanced lightweight multiwire cable to an Axona acquisition system (Axona Ltd., Herts, U.K., for the one microdrive-implanted animal). Both cables allowed the animal to move freely within the available space. Unit activity was amplified by a factor of 3000–5000 and band-pass-filtered 600–6000 Hz (Neuralynx) or from 800 –6700 Hz (Axona). Spike waveforms above a threshold set by the experimenter (50-80 μV) were time-stamped and digitized at 32 kHz (Neuralynx) or 48 kHz (Axona) for 1 ms. Local field potential (LFP) signals, 1 per tetrode for the hyperdrives and 1 in total per Axona microdrive, were amplified by a factor of 1000 and recorded continuously between 0 and 475 Hz at a sampling rate of 1893 Hz or 2000 Hz. In some Neuralynx recordings, the raw signals were also recorded (32 kHz). The LFP channels were recorded referenced to the ground screw positioned above the animal’s cerebellum (Neuralynx), or against an electrode from one microdrive tetrode (Axona). Light-emitting diodes (LEDs) on the headstage were used to track the animal's movements at a sampling rate of 25 Hz (Neuralynx) or 50 Hz (Axona).

### Behavioural procedures

The rats were food-restricted, maintaining their weight at a minimum of 90% of their free-feeding body weight, and were food deprived 12–18 hr before each training or recording session. During the 2–3 weeks between surgery and testing, the animals were trained to forage in an open field arena. The arena was a 150 x 150 cm square box with a black floor mat and 50 cm-high black walls, into which vanilla or chocolate biscuit crumbs were randomly scattered. Curtains were not used and abundant visual cues were available to the foraging rat. The animals were also habituated to a small sleep chamber, which was a 35 (l) x 40 (w) x 70 (h) cm plexiglas box with towels lining the floor and walls. This chamber was placed at the center of the larger open field arena, but due to the height and opacity of the sleep chamber’s walls, the animal was not able to see external cues. Between sessions in the open field and sleep chamber, the rat was placed next to the arena on an elevated flower pot lined with towels.

Each recording contained one or two open field foraging sessions and one or two sleep box sessions, with a combined duration of 20–95 minutes (median 43 minutes) for open field sessions and 55–360 minutes (median 97 minutes) for sleep box sessions. The open field arena floor was cleaned prior to beginning each recording. Data from multiple sessions of the same type was concatenated for analysis purposes. In some recordings, the animal also ran on a linear or circular track; the data from these sessions was not used in the present study. Intervals between successive sessions in a recording were of the minimum duration necessary to set up equipment for the following session (median 6 minutes). Recordings were generally performed during the dark and early-light phase of the 12h/12h light cycle.

### Spike sorting and single unit selection

Spike sorting was performed offline using a either manual or semiautomatic methods. MClust (A.D. Redish, http://redishlab.neuroscience.umn.edu/MClust/MClust.html) was used for manual sorting; semiautomatic sorting was done with KlustaKwik (K.D. Harris) followed by manual curation in MClust, or with Kilosort^70^ (https://github.com/cortex-lab/KiloSort) followed by manual curation in Phy (C. Rossant, https://github.com/kwikteam/phy). Spike rate autocorrelation and crosscorelation were used as additional tools for separating or merging spike clusters.

Single units were discarded if more than 0.5% of their interspike interval distribution was comprised of intervals less than 2 ms. Additionally, units were excluded if they had a mean spike rate of less than 0.5 Hz or greater than 10 Hz in either the sleep box or open field sessions.

### Spatial tuning curves

Animal position was estimated by tracking the LEDs on the headstage connected to the drive. Only time epochs where the animal was moving at a speed above 2 cm s^-1^ were used for spatial analyses.

To generate 2D rate maps for the open field arena, position estimates were binned into a 4 x 4 cm square grid. The spike rate in each position bin was calculated as the number of spikes recorded in the bin, divided by the time the animal spent in the bin. The resultant 2D rate map was smoothed with a gaussian kernel with σ = 1.5 bins.

The animal’s head direction was determined from the relative positions of LEDs on the headstage. Head-direction tuning curves were calculated by binning the head-direction estimates into 6-degree bins. The spike rate in each angular bin was calculated as the number of spikes recorded in the bin, divided by the time the animal spent in the bin. The resultant tuning curve was smoothed with a gaussian kernel with σ = 1 bin, with the ends of the tuning curve wrapped together.

### Identification and selection of grid cells

For all units, autocorrelations of the open field rate maps were generated as described previously^9^. From the autocorrelogram, a gridness score was calculated to quantify periodicity in the rate map. We modified a previously described method^23^, in order to improve its ability to detect strongly elliptical grid patterns.

Similar to previous methods, the algorithm iteratively defined an expanding annulus centred on the autocorrelogram’s origin. The annulus’ outer radius began at twice the radius of the central peak, and increased up to the width of the autocorrelogram. The annulus’ inner radius was always 80% of its outer radius. To define coordinates for sampling the autocorrelogram, lines were created radiating outward from the origin at equal angular intervals. For each line, the mean value of the autocorrelogram in the segment bounded by the annulus was calculated, yielding a circular “slice” of radial autocorrelogram samples.

To account for elliptical distortions of the grid pattern, we linearly transformed the annulus’ coordinates such that its bounding circles became elliptical. Specifically, the transformation mapped the coordinates of a unit circle onto the coordinates of an ellipse with orientation *θ*, semimajor axis *a* and semiminor axis *b*:

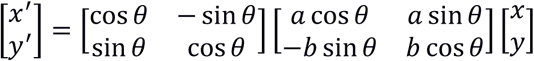

where the column vector 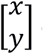 denotes a coordinate pair and 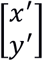 denotes the transformed coordinate pair. We iteratively adjusted the ellipse parameters and sampled the autocorrelogram at the transformed coordinates, thus accounting for many possible elliptical distortions of the autocorrelogram spanning the full 360° range of ellipse rotations and a range of eccentricities.

To compensate for the additional degrees of freedom introduced by the ellipse parameters, it was necessary to define more stringent criteria for identifying grid patterns in order to avoid detection of false-positives. Since grid patterns are characterized by both rotational and concentric periodicity, we defined two separate gridness scores: first, a “rotational” score was used to identify rotational symmetry (this has been the basis of previous gridness scores^9,54^), and second a “concentric” score which was used to identify concentric periodicity.

To calculate the rotational score, each elliptical slice of the autocorrelogram was compared with shifted versions of itself, corresponding to 30° rotations of the original circle. This was performed using unnormalized circular autocorrelation *R*_*yy*_, defined by:

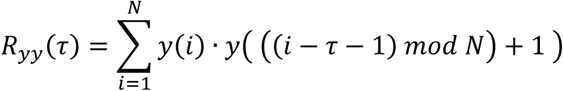

where *y* is the elliptical autocorrelogram slice, τ is the lag and *N* is the number of elements in the slice. The modulo operator wraps the ends of the slice together. According to the six-fold rotational symmetry of a grid pattern, the rotations were grouped as “in-phase” (60° and 120°) and “out-of-phase” (30°, 90° and 150°). The rotational gridness score was defined as the mean in-phase autocorrelation minus the mean out-of-phase autocorrelation.

The concentric gridness score was based on the principle that the autocorrelogram of a grid cell shows concentric periodicity, with maxima at each ring of peaks. To quantify this property, a second “inner” elliptical annulus was defined, having half the scale of the main “outer” elliptical annulus, and the same orientation. In the event of the outer elliptical annulus falling exactly over the inner ring of peaks in the autocorrelation, the inner elliptical annulus would fall along the valley between the central peak and the inner ring of peaks. The concentric gridness score was defined as the mean value of the autocorrelogram within the outer elliptical annulus, minus the mean value within the inner elliptical annulus.

The gridness scores for a cell were determined by finding the elliptical annulus that maximized the rotational gridness score. The same elliptical annulus was then used to determine the concentric gridness score. To establish a significance value, the scores were recalculated for shuffled experimental data. For each single unit, its spike train was time-shifted by a random amount of minimum 20 s and maximum 20 s less than the session’s duration, wrapping the end of the session to the beginning. 2D rate maps and autocorrelograms were generated from the shuffled data, and the rotational and concentric gridness scores were recalculated as described above. This process was repeated 1000 times for each unit, yielding a distribution of both rotational and concentric gridness scores. The shuffled rotational and concentric scores were individually z-score-normalized and then added together to yield a “combined” gridness score. The combined gridness score was then calculated for the original data, by applying the same transformation to its rotational and concentric scores as was applied to the shuffled data. Finally, the percentile of the cell’s combined gridness score was calculated with respect to the distribution of shuffled combined gridness scores. The unit was classified as a grid cell if its score exceeded the 99^th^ percentile of the shuffled distribution.

To ensure that phase relationships between grid cells could be reliably measured, additional constraints were applied to select grid cells for subsequent analysis. To select only cells with refined, regular grid patterns, cells were discarded if six inner autocorrelation peaks could not be identified within a small distance of the ellipse at which the optimal gridness score was evaluated, or if their open field rate map had a sparsity value greater than 0.5, where sparsity *s* is defined as:

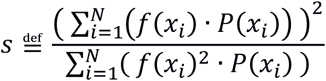

where *x*_i_ is the *i*^*th*^ position bin, *f*(*x*_i_) is the firing rate of the cell when the animal is located in the *i*^*th*^ position bin, and P(*x*_*i*_) is the probability of the animal being located in the *i*^*th*^ position bin.

Cells were also discarded if their grid spacing (defined as the mean distance of the six inner peaks from the origin) was greater than 1.4 metres.

### Grid module classification

According to a previously described method^32^, grid-cell module membership was classified on a pairwise basis: each cell pair was either defined as intramodular (both cells from the same module) or transmodular (cells from different modules). Briefly, an ellipse was fitted to the inner ring of peaks of each cell’s open-field rate map autocorrelogram (**Fig. 2a**). The area of the intersection *I* and union *U* of the two ellipses was calculated (**Fig. 2b**). The *I/U* ratio provided the module classification metric: cell pairs with a *I/U* ratio of 0.75 or greater were classified as intramodular, while cell pairs with a lower *I/U* ratio were classified as transmodular. While the method’s original description used a lower threshold (0.6), we found that almost all thresholds between 0.5 and 0.8 yielded significantly different distributions for intramodule and transmodule grid-cell pair correlations (**Supplementary Fig. 5a**).

### Grid phase offset measurement

Intramodular grid-cell phase (*ϕ*_*G*_) was quantified using a previously described method that improves the robustness of phase measurements to elliptical distortions of the grid pattern^32^. Briefly, the cross-correlogram of the two cells’ open-field rate maps was calculated, and the nearest peak to the origin was identified. An autocorrelogram was then generated from the cross-correlogram, and its inner Voronoi cell was used to define the boundary of the phase tile. *ϕ*_*G*_ was defined as the distance from the origin to the innermost peak, divided by the distance to the phase tile edge that was closest to the innermost peak.

### Identification of head-direction cells

For each cell’s head-direction (HD) tuning curve, its concentration parameter ? was estimated, and Rayleigh test for circular nonuniformity was performed. The cell was classified as a HD cell if the value of ? exceeded 1.0 and the Rayeigh *P*-value was less that 0.001. Cells that passed both the HD and grid criteria were classified as conjunctive grid-HD cells.

### Exclusion of duplicate recordings

Since the single unit recordings took place over successive days, with tetrodes being moved in small increments, many neurons may have been recorded on more than one occasion. Countermeasures were taken to minimize the probability of including duplicate recordings of the same cells or cell pairs. The positions of tetrodes were monitored across recordings, such that it was possible to localize where each unit was recorded. It has been reported that the majority of separable units in extracellular recordings are recorded within 50 μm of the soma^71,72^; therefore we classified units recorded within 50 μm of each other on the same tetrode as being potential duplicates.

Since the spatial firing patterns of grid and HD cells are highly stable across days, it was possible to use these characteristics to rule out many possible duplicate unit recordings. For grid cells and grid-HD cells, units recorded within 50 μm were compared by calculating the Pearson correlation coefficient of their open field rate maps. Cell comparisons with *r*-values below 0.5 were ruled out as duplicates. Similarly, for HD cells, possible duplicate recorded cells were ruled out if the Pearson correlation coefficient of their HD tuning curves was below 0.5.

For single and pairwise unit analyses, all units or unit pairs were judged to be unique according to the above criteria.

### Classification of sleep states

Periods of sleep were identified from when the animal was continuously immobile (speed < 1 cm s^-1^) for at least 120 s. To discriminate between slow-wave-sleep (SWS) and rapid-eye-movement (REM) sleep, the ratio between rhythmic activity in the theta (θ, 5–10 Hz) and delta (δ, 1–4 Hz) bands in the MEC LFP was used. The instantaneous amplitude in each frequency band was determined by calculating the absolute value of the Hilbert transform of the band-pass-filtered LFP. The instantaneous amplitude was smoothed with a gaussian kernel with σ = 5 s. Periods of sleep in which the value of θ/δ remained above 2.0 for at least 20 seconds were classified as REM; remaining sleep periods were classified as SWS.

### Analysis of pairwise spike rate coupling

Spike rates were calculated by binning spike times at a resolution of 10 ms and smoothing the resultant spike count vector with a Gaussian kernel with σ = 10 ms for SWS and σ = 150 ms for RUN and REM. The population rate was calculated by binning unsorted MEC multiunit activity at a resolution of 10 ms and smoothing with a Gaussian kernel with σ = 30 ms.

Two approaches were used for quantifying coupled spiking of cell pairs: the Pearson correlation coefficient, and a generalized linear model (GLM).

The Pearson method simply correlated the spike rate vectors of the two cells and used the resultant *r*-value as the metric for coupled spiking. Cross-correlations were calculated by iteratively lagging one of the spike rate vectors relative to the other.

The GLM framework is an established means for quantifying the contributions of multiple factors that may influence the spiking of a cell ^54,73,74^. We implemented a GLM to calculate the functional coupling between the spike rates of two cells A and B, while discounting common coupling to the global population spike rate, which is known to strongly influence pairwise spiking correlations^31^. The spike rate of cell A was modelled as an inhomogeneous Poisson process, as a function of both the spike rate of cell B and the population spike rate, using a log link function. MATLAB function “glmfit” was used to fit the model. To compensate for differences in the distributions of spike rates for single units and MUA spike trains, both regressors were z-score-normalized prior to fitting the model. The fitted model yielded a coefficient for the coupling between the spike rates of cells A and B (here referred to as *β*), as well as a coefficient for the coupling of cell A to the population spike rate (here referred to as *β*_POP_). As an analog to conventional cross-correlation, the GLM spike rate cross-correlation (SRC) was defined, for a given time lag τ, as the *β*-value yielded by fitting the GLM after adding τ to the spike times of cell A.

### Random-walk modelling of grid and HD spiking correlations

We refer a previously employed rationale^29^, that for two cells whose spiking is tuned to a variable *x* expressing random-walk dynamics (such as an animal’s head direction or position), the temporal spike rate cross-correlogram (SRC) of the two cells is determined by the velocity distribution of *x*.

Suppose that the spike rates of two neurons are tuned to one-dimensional random-walk process *x* with tuning curves *f*_1_(*x*) and *f*_2_(*x*), giving rise to spike trains with rates *s*_1_(*t*) and *s*_2_(*t*), where *t* represents time.

The unnormalized cross-correlation between the spike rates of two neurons *R*_*s*_1_,*s*_2__ is defined by:

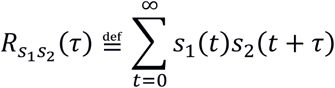

where *τ* is the time lag. If *v* represents the velocity of *x*, then at a particular time lag τ, the value of *x* changes by amount *v*τ. Therefore, for a particular value of *v*, the SRC can be approximated by the correlation of the two neurons’ tuning curves, offset by *vτ*:

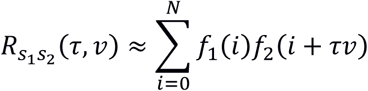

where *N* is the number of bins in the tuning curve and *i* is the tuning curve bin index. If we assume that *v* is a random variable with a known distribution, a more general approximation can be obtained by averaging across all possible velocities, indicated here by ⟨·⟩_*v.*_

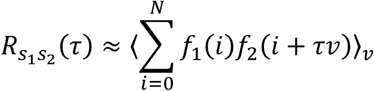

Thus, the distribution of *v* determines the temporal scaling of the cross-correlatione of *s*_1_ and *s*_2_.

To estimate the average speed of the grid position variable underlying grid-cell SRCs, we modelled the variable as a 2-dimensional random walk process with a specified mean speed. At each step of the random walk, the direction of the discplacement vector was randomly drawn from a uniform distribution, while the vector’s magnitude was drawn from a chi-squared distribution with two degrees of freedom, multiplicatively scaled such that its mean matched the desired value.

For each pair of grid cells, we fitted a parameterized grid pattern to their rate maps which preserved the phase offset and mean spike rate of the cell pair. Spike trains were simulated for each of the two cells as inhomogeneous poisson processes with rates determined by sampling its grid pattern at the coordinates of the random walk. The GLM SRC was then calculated from the simulated spike trains. This process was repeated using a wide range of random walk speeds, and the speed that yielded the GLM spike rate cross-correlation that most closely matched the original was selected as the estimated speed. For finding the optimal match, we used a cost function that used equally weighted contributions from the Pearson correlation coefficient and the mean squared difference between the simulated and original cross-correlations. Some cell pairs (particularly those with intermediate phase offsets) had empirical SRCs that were relatively flat, which yielded unreliable model fits; we therefore excluded cell pairs whose optimal model fits were poor (specifically, if their Pearson correlation with the original cross-correlation was less than 0.25).

For HD cells, the speed of the head direction signal was estimated as for grid cells, except that a 1- dimensional angular random walk was used instead. The spike trains of HD cells were simulated as inhomogeneous poisson processes whose rates were sampled from the cells’ HD tuning curves at the coordinates of the random walk.

### Detection of CA1 sharp-wave-ripple (CA1-SWR) events

A previously described method^75^ was followed. Briefly, the hippocampal local field potential signal was band-pass-filtered between 125-250 Hz and rectified, and the instantaneous amplitude was quantified by fitting a cubic spline to the peaks of the rectified filtered signal. CA1-SWR events were identified by detecting time periods where the 125–250 Hz amplitude surpassed a threshold of 3.5 standard deviations from the mean. A lower threshold of 2.0 standard deviations from the mean was used for defining the start and end times of each CA1-SWR event.

### Detection of MEC sharp-wave (MEC-SW) events

The MEC-SW is a negative deflection in the MEC LFP occuring most prominently in layer III, immediately following a CA1-SWR^49,50^. In each recording, one tetrode was designated for detecting MEC-SW, on the basis of visually identified sharp-wave events being present in the LFP. To detect MEC-SW, the MEC LFP from SWS epochs was band-pass-filtered between 10–40 Hz. Negative peaks with an amplitude exceeding 4 standard deviations from the mean were classified as MEC-SW events. Only SWS epochs were considered for identifying MEC-SW events, to avoid possible interference from harmonics of the theta oscillation, which fall in the same frequency range of MEC- SW.

### Histology and reconstruction of tetrode placement

Rats received an overdose of sodium pentobarbital and were perfused intracardially with saline followed by 4% formaldehyde. The brains were extracted and stored in 4% formaldehyde, and frozen sagittal sections (30 μm) were cut and stained with cresyl violet (Nissl). Each section through the relevant part of MEC /PaS was collected for analysis. All tetrodes from the 14-tetrode bundle were identified from digital photomicrographs by comparing tetrode traces from successive sections.

### CA1 dataset

The recordings of CA1 putative pyramidal cells were obtained from a previously published dataset^43^, downloaded from http://dx.doi.org/10.6080/K0862DC5. Only data from the “PRE” rest session was used, and the original arousal state labels were used. From 3 animals in which large numbers of pyramidal cells were recorded on both probes, a total of 6 recordings were used. In each recording, a random sample of 200 pyramidal cell pairs was taken. Pairs were exclusively formed of cells on separate probes, to prevent spiking correlations being spuriously affected by erroneous spike sorting or missed detections of temporally overlapping spikes. Cells were excluded if their mean spike rate was below 0.2 Hz during any of the four 15-minute divisions of the first hour of SWS. Population spike rates were quantified by combining activity from all pyramidal cells.

### Data analysis and statistics

Data analyses were performed with custom-written scripts in MATLAB (MathWorks). All statistical tests were two-tailed. Each test used was judged to be the most suitable for the data in question. Where possible, nonparametric methods were used in preference to parametric methods, to avoid making unwarranted assumptions about the underlying data distributions. All confidence intervals were estimated using bootstrap resampling (*n* = 1000 resamples). Spearman rank was used for all correlation measurements apart from for cell-pair spike rate correlations, where Pearson correlation was used, following convention. Comparisons which lacked established testing methods were performed using nonparametric resampling methods. Samples included all available cells that matched the classification criteria for the relevant cell type, but excluded repeated recordings of the same cells (see “exclusion of duplicate recordings”). The study did not involve any experimental subject groups, therfore random allocation and experimenter blinding did not apply and were not performed.

### Code availability

Code is available on request from the authors.

## Acknowledgments

We thank A. Z. Vollan for assistance with recordings and A.M. Amundsgård, K. Haugen, E.Kråkvik, and H. Waade for various technical assistance. The work was supported by an Advanced Investigator Grant from the European Research Council (GRIDCODE – grant no. 338865), the European Commission’s FP7 FET Proactive Programme on Neuro-Bio-Inspired Systems (GRIDMAP - Grant Agreement 600725), a NEVRONOR grant from the Research Council of Norway (grant no. 226003), the Centre of Excellence scheme and the National Infrastructure Scheme of the Research Council of Norway (Centre for Neural Computation, grant number 223262; NORBRAIN1, grant number 197467), the Louis Jeantet Prize, the Körber Prize, and the Kavli Foundation.

## Supplementary figure legends

**Supplementary Figure 1:** Basic LFP and single-unit spiking characteristics

**a:** Example traces of MEC activity during RUN, SWS and REM. From top to bottom: combined spike rate of all recorded units; spike rasters for 8 grid cells; wideband LFP signal. **b:** MEC LFP power spectra generated from the recording shown in (a). The power spectra were calculated in 5-second windows using the multitaper method, using all available time periods for each state. The lines and shaded areas represent mean ±S.E.M. **c:** Distributions of mean spike rates of all units during RUN, SWS and REM (*n* = 138 grid cells, 95 HD cells, 39 grid-HD cells). Box plots show the median, interquartile and full range of mean spike rates in each group. **d:** Comparison of mean spike rates of grid, HD and grid-HD cells between RUN, SWS and REM (cell types are displayed in the same vertical order as in (c)).

**Supplementary Figure 2:** Population coupling effects on Pearson and GLM correlation measures

**a:** Demonstration of effect of global population rate modulation on correlations. **a1** shows spike trains from two simulated neurons producing inhomogeneous Poisson spike trains (black rasters, top) with rates oppositely modulated by a signal of interest (grey line). The Pearson and GLM spike rate cross-correlograms (SRCs) of the spike trains (bottom) show the time course of the relationship between the two neurons’ spike rates. **a2** shows the same situation as (a1), but with the two neurons’ spiking also positively modulated by global population rate fluctuations (red line). The population rate strongly impacts the Pearson SRC, whose zero-lag deflection changes from negative to positive. Conversely, the GLM SRC retains a similar shape to the SRC in (a1). **b:** Relationship between zero-lag Pearson (*r*-value) and zero-lag GLM coupling coefficient (*β*_0_) for grid/grid (top) and HD/HD (bottom) cell pairs. Note that cell pairs of both types with Pearson *r*-values near zero are generally negative when measured with the GLM method. **c:** Coupling of single unit spiking to population spike rate. A Poisson GLM was fitted to the spike rate of each unit, with the population spike rate as the single regressor. The plots show the resultant distributions of zero-lag GLM couplings for the population rate (*β*_POP_). Population rates were z-score transformed before fitting the model, such that results were comparable across different mean population rates. Note the almost exclusively positive *β*_POP_-values.

**Supplementary Figure 3:** Spiking correlations between pairs of grid cells

**a:** Scatter plots showing the lag time of the largest peak for each GLM SRC, versus the corresponding GLM *β*_0_. For cell pairs with positive *β*_0_, the largest positive peak was identified; for cell pairs with negative *β*_0_, the largest negative peak was identified. Most of the peaks falling far from zero lag are for cell pairs with small absolute *β*_0_, indicating weak correlations. **b:** Histograms of the peak times shown in (a). **c:** Histograms of *β*_0_ distributions (*n* = 138 grid, 95 HD, 39 grid-HD). **d:** Relationship between the grid phase offset *ϕ*_*G*_ of grid/grid cell pairs and their zero-lag correlation, measured with the Pearson correlation coefficient. Note the lack of strong negative correlations predominance of negative correlations, in contrast to the strong correlations evident when using the GLM approach in **Fig. 1c**. **e**: Relationship between grid/grid cell pair rate map similarity (measured as the *r*-value of the Pearson correlation between each cell pair’s respective rate maps), and GLM *β*_0_.

**Supplementary Figure 4:** Signatures of hexagonal geometry in grid-cell-pair spiking correlations

**a:** A simulated triangular grid pattern. Each grey dot represents a vertex (receptive field) of the grid. The hexagonal tiles surrounding the vertices are Voronoi cells which form the phase space of the grid pattern. The black arrow marked “*s*” denotes the distance between adjacent vertices (the grid spacing). Within the central tile are drawn a series of concentric rings, indicating zones corresponding to different values of *ϕ*_*G*_, the magnitude of a grid phase offset vector. Note that the phase tile border encroaches on the outermost *ϕ* zone, meaning that the largest values of *ϕ*_*G*_ have a nonuniform radial distribution. **b:** Illustration of the four spatial kernel types used to simulate grid firing fields. **c:** Rate map similarity (RMS, defined as the Pearson correlation coefficient of the two grid patterns’ simulated spike rate maps) scores for simulated pairs of grids as a function of phase offset. Each position on the hexagonal phase tile represents a particular phase offset of one grid from the other. The colour at each point indicates the RMS value of a grid-cell pair of that phase offset. Each plot shows the result of simulations using a different spatial kernel type. In each case, the normalized field width parameter *w* was 0.19, equal to the empirically estimated value. Contour lines follow equal RMS values. Note the white ring where RMS is zero, and the non-concentric deviations near the phase tile’s edge. Inset: examples of simulated rate maps. **d:** Relationship of RMS with *ϕ*_*G*_ and field width *w*. The dashed line traces the path of the value of *ϕ*_*G*_ at which RMS crosses zero (*ϕ*_*G*0_) Note the convergence of *ϕ*_*G*0_ on a value of 0.33 as *w* increases. The black circle on the first plot indicates *ϕ*_*G*0_ at the empirically estimated value of *w* for the sample of grid cells (see f,g). **e:** Empirical relationship between *ϕ*_*G*_ and RMS (black dots) among grid/grid pairs. The grey line indicates the values obtained from simulated rate maps for grid/grid pairs with uniform-randomly distributed phases. **f:** F1 scores for dividing the distribution of grid/grid pair zero-lag GLM *β*-values into positive and negative values at different *ϕ*_*G*0_-values. The vertical dotted grey line indicates the empirical *ϕ*_*G*0_- value of 0.32. **g:** *ϕ*_*G*_-values that maximise the F1 score as shown in (f), with 95% bootstrap confidence intervals. The dotted line indicates the simulated *ϕ*_*G*0_-value of 0.32. **h:** Relationship between grid spacing and *w*.

**Supplementary Figure 5:** Effect of module classification threshold on zero-lag correlation comparisons. When classifying grid-cell pairs as intramodular or transmodular, a threshold *I/U* ratio of 0.75 was used. To ensure that the outcomes shown in **Fig. 2f** were not dependent on the particular threshold value used, we examined the results of using different threshold values.

**a**: Relationship of *I/U* ratio module classification threshold on the two-sample Kolmogorov-Smirnov *P*-value when comparing the cumulative distributions of zero-lag correlations (*β*_0_) for intramodular and transmodular pairs, as shown in **Fig. 2g**. Each line shows the resultant *P*-values for each threshold in a given arousal state. The bars at the top of the plot indicate the threshold values for which *P* < 0.05. The dashed black line indicates the actual threshold that was used in analyses (0.75). **b**: Same as (a), but showing *P*-values for the 2-sample Student’s t-test.

**Supplementary Figure 6:** Spike rate correlations between grid and conjunctive grid-HD cells

**a**: Colour-mapped GLM SRCs for all intramodular grid/grid-HD pairs, ranked by their spatial phase offset magnitude *ϕ*_*G*_. Each row of the matrix is the SRC of one cell pair. The black line to the right of the matrix indicates the value of *ϕ*_*G*_ for each pair. **b:** Relationship between grid phase magnitude *ϕ*_*G*_ and GLM zero-lag coupling (*β*_0_) during each state. Displayed statistics are for Spearman rank correlation (*n* = 16 grid/grid-HD pairs). **c:** Same as (a), but with cell pairs ranked in order of the *β*_0_- value during RUN. **d:** Same as (b), but showing relationship between *β*_0_ during RUN and *β*_0_ during SWS /REM.

**Supplementary Figure 7**: Stability of spike rate coupling in MEC and CA1 cell pairs

**a:** Same as **Fig. 5a**, with subsampled spike rates. Since mean spike rates influence the magnitude of observed correlations^53^, it is possible that differences in observed correlation stability could merely reflect differences in mean spike rate. To discount this possibility, we randomly subsampled each unit’s mean spike rate to 0.2 Hz during each 15-minute window of the first hour of SWS. *β*_0_-values are compared between the first 15 (*β*_0_0-15_) and last 15 minutes (*β*_0_45-60_) of the first hour of SWS. Displayed *r*-values are for Spearman rank correlation. **b:** Stability of zero-lag spike rate correlations for different Gaussian kernel widths, for intramodular grid/grid, HD/HD and CA1/CA1 cell pairs, calculated as the correlation between the cell pairs’ *β*_0_0-15_ and *β*_0_45-60_ values. **c:** Coupling stability for CA1 putative pyramidal cell pairs within each recording. Each plot shows, for one recording, all CA1 pyramidal cell pair GLM spiking correlations from the first 15 (*β*_0_0-15_) and last 15 minutes (*β*_0_45-60_) of the first hour of SWS.

